# Gene-sized DNA insertion at genomic safe harbors in human cells using a site-directed transposase

**DOI:** 10.1101/2025.09.16.676703

**Authors:** James E. Short, Lisa Sharek, Joshua F. Meckler, Ilko Stoytchev, Christopher T. Tran, Clara Errard, Brian E. Hew, Brandon E. Johnson, David F. Waller, Shanfu Xie, Joseph J. Higgins, Raymond Tabibiazar, Jesse B. Owens

## Abstract

Achieving precise and efficient integration of gene-sized DNA sequences into the human genome remains a major obstacle to gene therapy. Existing approaches depend on double-strand DNA breaks, which can lead to unintended genome alterations. Many monogenic diseases arise from diverse patient-specific mutations, making individualized correction impractical and underscoring the need for universal full-gene replacement strategies. We developed INsertion by Targeted Anchoring and Conditional Transposition (INTACT) to enable targeted insertion at genomic safe harbor loci. We engineered a mammalian transposase with mutations in its DNA-binding domain to reduce off-target integration. Site specificity was then restored by linking programmable sequence-specific DNA-binding proteins to the transposase. Systematic optimization of INTACT revealed key determinants of precision, including non-covalent linkage between the transposase and DNA-binding protein, strict spacing between the binding site and the TTAA insertion sequence, and linkage of the DNA-binding protein to an internal position within the transposase. On-target insertion was achieved across multiple loci, with optimized INTACT averaging 1.2 targeted insertions per cell. An off-target assay confirmed that DNA-binding domain mutations substantially reduced unwanted integration events to near-background levels. Our site-directed transposase enables precise, efficient genomic insertion of >4kb DNA without double-strand breaks, offering a powerful new tool for genome engineering.

## INTRODUCTION

Integrating gene-sized DNA sequences into precise locations within the human genome remains a significant challenge in gene therapy (1,2). Many genetic conditions arise from distinct patient-specific mutations, necessitating customized genome editors for each case (1). A single therapy capable of inserting large DNA sequences to replace the entire defective gene could theoretically treat these conditions regardless of the specific mutation. Although virus-based methods are highly efficient, they lack control over the insertion site, raising concerns about potential adverse effects, such as insertional mutagenesis and oncogenesis (3–7). Moreover, non-integrating viral vectors such as AAV can be immunogenic, making repeat dosing problematic (8–11). Although RNA-guided CRISPR-Cas systems can employ homology-directed repair (HDR) for site-specific insertion, this mechanism is inefficient in non-dividing cells and is often impractical for gene-sized DNA sequences (12,13). Alternate approaches such as homology-independent targeted integration (HITI) have been developed but, like all nuclease-based approaches, they introduce double-strand breaks (DSBs) that can lead to undesirable consequences like deletions, chromosomal rearrangements, and chromothripsis through the activation of non-homologous end joining (NHEJ) pathways (14–18). Genomic cleavage by CRISPR nucleases has been shown to activate a DNA damage response by the p53 tumor suppressor, resulting in unwanted selection for cells with impaired p53 function (19,20). Furthermore, CRISPR-based tools such as base and prime editors, which do not require DSBs, are limited by the availability of protospacer adjacent motifs (PAMs) and are restricted in the size of the edits they can introduce (21,22).

The discovery of transposable elements by Barbara McClintock over 75 years ago laid the groundwork for the development of gene therapy platforms based on these elements (23). Transposases are capable of integrating DNA sequences into the genome without the need for DSBs, presenting a safer alternative to nuclease-based methods (24). Among the transposable elements being explored for human gene therapy are *piggyBac*, *Sleeping Beauty*, and the only known active transposable element isolated from a mammal (the little brown bat), *Myotis lucifugus* transposase (MLT, or *piggyBat*) (25–29). These mobile elements use a single transposase enzyme to integrate large DNA sequences (as many as 100 kilobases) at TA dinucleotides (*Sleeping Beauty*) or TTAA tetranucleotides (*piggyBac*, MLT) (28,30). The DNA cargo is flanked by left and right transposon ends (also called terminal inverted repeats (TIRs)) that are excised by the transposase, forming a transpososome that inserts the transposon DNA without generating DSBs (24,31).

A transposable element system capable of precisely integrating large, gene-sized DNA sequences at defined genomic sites would be a powerful tool for gene therapy. In our previous work, we demonstrated that fusing DNA-binding proteins such as transcription activator-like effectors (TALEs) or catalytically inactive ‘dead’ Cas9 (dCas9) to the *piggyBac* transposase could facilitate targeted DNA insertion, detected in clonally expanded cells with single insertions. (32,33). However, these approaches required screening from polyclonal populations due to low targeting efficiencies (∼0.06%). Other attempts to link DNA-binding proteins to transposases have similarly produced low efficiencies (34–44). Recently, CRISPR-associated transposase (CAST) systems have been developed, demonstrating high efficiency RNA-guided targeting in bacterial cells (45,46). However, initial efforts to apply CAST systems in human cells had limited success, with targeting efficiencies of 0.6% in unsorted and 4.7% in sorted cells measured by droplet digital PCR (ddPCR) (47). Additionally, the therapeutic use of CAST is hindered by its reliance on a complex mix of components (a guide RNA plus eight different proteins), including the bacterial accessory factor ClpX, which is toxic to human cells (47).

To design a programmable transposase, we used the AlphaFold2 predicted structure to identify a set of mutations that disrupted the native DNA-binding function of MLT and prevented integration but preserved its catalytic activity for excision (Exc⁺Int⁻ mutants), a strategy previously devised for *piggyBac* (32,39,48). We next restored the transposase’s ability to mediate DNA integration by tethering it to DNA-binding proteins that anchor the complex to defined genomic loci. This engineered system, which we term INsertion by Targeted Anchoring and Conditional Transposition (INTACT), enables targeted integration at user-defined sequences.

We systematically evaluated several factors influencing INTACT, including: (i) spacing between the DNA-binding protein’s target sequence and the TTAA insertion site, (ii) target sequence orientation relative to the TTAA site, (iii) whether one or two DNA-binding protein target sequences flank the TTAA site, (iv) transposase linkage site (N-terminus, C-terminus, or internally), (v) DNA-binding protein type (zinc finger or TALE), (vi) surrounding sequence adjacent to the TTAA site, (vii) location of the genomic target sequence, (viii) whether the binding protein was covalently or non-covalently linked, and (ix) mutations in the transposase DNA-binding domain to reduce off-target insertion while maintaining efficiency.

Using a plasmid-based assay, we identified strict targeting rules for INTACT. Key factors included a non-covalent transposase-DNA-binding protein linkage and strict base-pair spacing between the binding protein’s target sequence and TTAA site. Applying these rules to safe harbor sites, we achieved high-efficiency on-target insertions at multiple loci. The most effective configuration targeted multi-copy sites, resulting in most cells receiving at least one targeted insertion, with an average of 1.2 on-target insertions per cell. Off-target analysis revealed near-background activity, indicating that DNA-binding domain mutations successfully reduced unintended integrations. This programmable transposase system enables site-specific integration without inducing DSBs and minimizes off-target integration, offering a safe and efficient method for inserting gene-sized DNA into the human genome.

## METHODS

### Plasmid construction

Plasmids were generated via Gibson cloning using the NEBuilder HiFi DNA Assembly reagent (New England Biolabs). Plasmid DNA for transfection was purified by ZymoPURE Plasmid Miniprep Kit or ZymoPURE Express Plasmid Midiprep Kit (Zymo). The primary plasmids used in this study can be obtained from Addgene. Descriptions of all 123 plasmids are included in Supplementary Table 1 and maps are available upon request.

#### Transposase (helper) plasmids

To generate the MLT helper plasmid, the gene for the full-length *Myotis lucifugus* transposase (MLT) was codon-optimized for human expression and synthesized as a gBlock gene fragment (Integrated DNA Technologies) and cloned into pcDNA3.3-TOPO (25). For covalent approaches including E2C MLT and HoD MLT plasmids, zinc finger (ZnF) or TALE DNA binding proteins were synthesized as gBlock gene fragments and cloned after the G66 position within the N-terminal domain. The DNA binding proteins contained a GGGS linker at their N-terminus and a GGGGS linker at their C-terminus. To generate ALFA MLT, insertion of the ALFA tag (49) at position G66 was performed using primers containing the ALFA tag sequence (TCCAGACTGGAAGAGGAACTGAGAAGAAGGCTCACAGAA) and Gibson cloning. Plasmids expressing fusion domains containing the nanobody that binds the ALFA tag (NbALFA) and including ZnF or TALE DNA binding proteins were synthesized as gBlock gene fragments and cloned into pcDNA3.3-TOPO. All TALEs used an N-terminal domain allowing for the recognition sequence to start with any base (50).

#### Excision plasmids

To construct the excision assay reporter plasmid, a gBlock was synthesized containing the ZsGreen gene, which was disrupted at a naturally occurring TTAA site by a 607 bp transposon. This transposon included full-length MLT left and right transposon ends flanking a polyadenylation signal (25). The disrupted gene was placed under the CAG (cytomegalovirus (CMV) immediate early enhancer, chicken b-actin promoter and b-globin intron) promoter. Upon transposon excision, seamless repair restores the TTAA site within the ZsGreen gene, restoring the uninterrupted sequence.

#### Integration plasmids

To construct the integration assay donor plasmid, the *piggyBac* transposon end sequences from the pB GIN donor (32) were swapped with MLT left and right transposon end sequences that were synthesized as gBlocks and cloned by Gibson assembly (25).

#### Plasmid-to-plasmid (P2P) reporter assay components

Donor and reporter plasmids for the P2P assay were modified from the previously described EmGFP splicing based reconstitution assay (51). To prepare the P2P donor plasmid, the 5’-half of EmGFP and SMN1 intron was PCR amplified from 1/2 EmGFP SMN1 transposon and placed under the CMV promoter. The donor transposon was flanked by the transposon end sequences of MLT that were synthesized as gBlocks. To construct the reporter plasmid, the 3’-half of EmGFP was PCR amplified from pT4 SMN1 2/2 EmGFP and cloned downstream of a 639 bp target region containing 26 randomly spaced TTAA sites. Reporter plasmids contained a unique DNA binder recognition sequence at the center of the target region. The ‘e2c’ DNA recognition sequence is bound by the E2C ZnF. Reporter plasmids, 1x e2c and 2x e2c contained a single or two e2c sequences placed 15 bp away from the target TTAA, respectively. The control reporter plasmid (no e2c) did not contain a binder recognition sequence. Reporters containing endogenous sequences contained 86 bp of genomic sequence inserted into the target region. For experiments interrogating the required spacing of recognition sequences to the TTAA, all other TTAA sites within 100 bp of the single target TTAA were removed and e2c sequences were inserted flanking the target TTAA at the indicated spacing on either side. Control conditions without functional helper plasmid used the pcDNA3.3 backbone, designated as ‘stuffer’.

### General culture conditions

HEK293T cells (Thermo Fisher Scientific NC0260915) were maintained in Dulbecco’s Modified Eagle Medium (DMEM) with GlutaMAX (Gibco) supplemented with 10% heat inactivated fetal bovine serum FBS (Neuromics), 1x Penicillin/Streptomycin (Gibco), and 0.75 µg/mL Amphotericin B (Gibco). Cells were grown at 37°C with 5% carbon dioxide (CO2) and passaged twice per week.

### Excision assay

The excision assay used a ZsGreen fluorescent protein gene disrupted by an MLT transposon. Transposase expression resulted in seamless excision, restoring ZsGreen expression, that was quantified by flow cytometry. HEK293T cells were seeded in 12-well plates at a density of 4.8 x 10^5^ cells per well. The following day, cells were co-transfected with the excision assay reporter plasmid and a MLT helper plasmid at a 3:1 ratio (900 ng of reporter plasmid: 300 ng of helper plasmid). Transfections were performed using the X-tremeGENE9 DNA transfection reagent (Roche) at a 1:3 ratio (1.2 µg of DNA: 3.6 µl transfection reagent per well). Following four days of culture, the percentage of ZsGreen+ cells was measured by flow cytometry using an Attune NXT flow cytometer (Thermo Fisher Scientific).

### Integration assay

The integration assay used a donor plasmid with a transposon containing a constitutively expressed TurboGFP fluorescent protein gene. Co-transfection with a MLT transposase helper plasmid resulted in transposon integration into the genome. Following 17 days of culture, stably expressing cells were measured by flow cytometry. Similar to the excision assay, HEK293T cells were seeded in 12-well plates. The following day, cells were co-transfected with the integration assay donor plasmid and a MLT helper plasmid at a 1:1 ratio (600 ng of donor plasmid: 600 ng of helper plasmid). Transfections were performed using the X-tremeGENE9 DNA transfection reagent (Roche) at a 1:3 ratio (1.2 µg of DNA: 3.6 µl transfection reagent per well). Following two days of culture, half the cells were resuspended and the percentage of TurboGFP+ cells was measured by flow cytometry to assess transient transfection efficiency. The remaining cells were cultured until 17 days post transfection, and the percentage of TurboGFP+ cells were again measured by flow cytometry. To normalize for transfection efficiency, the final integration efficiency was calculated by dividing the percentage of GFP+ cells from day 17 by day 2.

### Plasmid to plasmid reporter (P2P) assay

The P2P assay measured site-directed plasmid targeting. A donor plasmid carried a transposon with a CMV promoter driving the 5’ half of EmGFP, followed by a splice donor sequence. A reporter plasmid contained the target sequence, a splice acceptor, and the 3’ EmGFP fragment. Transposon insertion near the target sequence restored EmGFP expression, that was quantified by flow cytometry. PCR amplification of transposon-reporter junctions identified insertion sites. HEK293T cells were seeded in 24-well plates at a density of 2.4 x 10^5^ cells per well. The following day, cells were co-transfected with 300 ng of helper plasmid, 150 ng of P2P donor plasmid and 150 ng of reporter plasmid. Transfections were performed using TransIT-2020 (Mirus Bio) at a 1:3 ratio (0.6 µg of DNA: 1.8 µl transfection reagent per well). Following three days of culture, half the cells were pelleted and the remaining cells were measured by flow cytometry to detect the percentage of EmGFP+ cells. To prepare PCR template, cell pellets were lysed with 90 µl DirectPCR Cell lysis reagent (Viagen) with 1.0 mg/ml proteinase K (Meridian Bioscience) and rotated in a hybridization oven for three hours at 55°C, then heat-inactivated for 45 min at 85°C. PCR was performed using 1 µl of lysate, 10 µl of KOD ONE BLUE polymerase 2x mastermix (Toyobo), 7.8 µl water and 0.6 µl each primer (10 µM). The forward primer was located on the reporter plasmid and the reverse primer was located on the donor transposon. PCR cycle conditions were 95°C for 30 sec followed by 32 cycles of 98°C for 10 sec, 60°C for 5 sec, and 68°C for 10 sec. Products were run on a 2% agarose gel in Tris-acetate-EDTA buffer and fragments for Sanger sequencing were extracted using the Zymo Gel DNA recovery Kit (Zymo).

### Safe harbor genome targeting

The donor plasmid was co-transfected with helper plasmids encoding engineered MLT, guided to the genome by DNA-binding proteins specific to the described frameworks. Insertions of the transposon at genomic target sites were detected by ddPCR using genomic DNA as template. HEK293T cells were seeded in 24-well plates at a density of 2.4 x 10^5^ cells per well and the following day, unless indicated, cells were co-transfected with 100 ng of helper plasmid and 500 ng of the same donor plasmid used in the P2P assay. Transfections were performed using TransIT-2020 (Mirus Bio) at a 1:3 ratio (0.6 µg of DNA: 1.8 µl transfection reagent per well). Following three days of culture, the cells were pelleted and lysed to prepare genomic DNA for ddPCR, as described above (90 µl DirectPCR Cell lysis reagent (Viagen) with 1.0 mg/ml proteinase K (Meridian Bioscience) was rotated for three hours at 55°C then heat-inactivated for 45 min at 85°C). To reduce the influence of variable transfection efficiency, HEK293T cells in the experiment shown in Fig. 7d were sorted for expression of the near-infrared fluorescent protein iRFP670 to ensure uptake of the DNA. HEK293T cells were seeded in 6-well plates at a density of 9.6 x 10^5^ cells per well. Using the final optimal ratio that was determined in the experiment shown in Fig. 7c, cells were co-transfected with 280 ng of ALFA MLT m12 helper plasmid or indicated control helper plasmid, 60 ng of each of rDNA1 NbALFA TALE fusion plasmids, 1980 ng of donor plasmid, and 20 ng of piRFP670-N1 plasmid (Addgene), which constitutively expresses iRFP670, and cultured for 4 days. A FACSAria IIu flow cytometer (BD Biosciences) was used to sort the brightest ∼20% of piRFP670+ live cells (excitation 635 nm, emission 780/60 nm).

Following FACS, 50 µl of cells in Live Cell Imaging Solution (Thermo Fisher Scientific) were combined with 50 µl of DirectPCR Cell lysis reagent (Viagen) with 1.0 mg/ml proteinase K (Meridian Bioscience) and rotated for three hours at 55°C, then heat-inactivated for 45 min at 85°C to prepare for ddPCR analysis.

### Droplet digital PCR (ddPCR)

ddPCR was performed on cell lysates from safe harbor genome targeting experiments to determine the frequency of targeted integrations. Reactions (22 µl) contained 0.5 µl of cell lysate, 11 µl of 2X ddPCR Supermix for Probes (BioRad), 10 units XhoI (New England Biolabs), target and reference primers at 900 nM and probes at 250 nM. The plate was sealed with a PX1 PCR Plate Sealer (BioRad). Droplets were prepared using the QX200 AutoDG Droplet Generator System (BioRad). PCR cycle conditions were 95°C for 10 min followed by 45 cycles of 95°C for 30 sec, 60°C for two min and 72°C for two min, and a final incubation at 98°C for 10 min. Droplets were read on a QX200 Droplet Reader (BioRad) and analyzed using QX Manager 1.2 Standard Edition software (BioRad). The number of on-target insertions per 100 cells was calculated by multiplying the ratio value of target to reference by the number of reference copies per cell. The UBE2D2 reference sequence, located on chromosome 5, was previously shown to be present in three copies per cell in HEK293T cells (52). We confirmed this copy number by comparing the UBE2D2 reference to single and double copy inserts originating from prime editing performed in HEK293T single cell clones (53). Because HEK293T cells have abnormal karyotypes with variable chromosome copy numbers, the choice of reference locus can significantly affect the calculated ratio of target to reference and the resulting efficiency per ‘genome’. For example, a reference on a disomic chromosome would yield a higher calculated efficiency than one on a trisomic chromosome. To avoid over-or under-reporting due to user reference selection, we report total integration efficiency per cell. A description of all primers and probes is found in Supplementary Table 2.

### Identification of preferred octanucleotide sites by whole genome insertion site recovery using the self-reporting transposon (SRT) assay

The SRT protocol was performed as previously described (54). Briefly, the SRT donor plasmid contained a constitutively expressed hygromycin B gene lacking a polyadenylation signal. RNA polymerase II transcription extended beyond the hygromycin B gene, through the transposon end, and into the flanking genomic sequence. Sequencing of SRT-derived transcripts identified transposon-genome junctions, enabling mapping of insertion sites. HEK293T cells were seeded in 24-well plates and the following day, cells were co-transfected with 100 ng of helper plasmid encoding unmodified MLT and 500 ng of SRT donor plasmid. Cells were cultured under hygromycin selection for 3 weeks and pelleted for SRT analysis. RNA was isolated using the RNeasy Plus Mini Kit (Qiagen) from hygromycin B selected cells. RNA was reverse transcribed to cDNA using Maxima H Minus Reverse Transcriptase (Thermo Fisher Scientific). Self-reporting transcripts were generated by PCR-amplification from cDNA template. Libraries were prepared using the Nextera XT DNA Library Preparation Kit (Illumina) and were sequenced on a NextSeq 2000 (Illumina).

### Bioinformatics

#### Identification of SRT genomic insertion sites and flanking sequence patterns

For the SRT pipeline, we employed a modified bioinformatic algorithm as described previously (54). For each sample, we demultiplexed the 150 x 2 reads using i7 and i5 barcodes, where i5 was inline in read1 (R1). Cutadapt (55) was used for trimming the reads, removing adapters, and extracting the inline i5 barcode and the left transposon end sequence. The TTAA was retained during trimming. The R1 sequences were aligned to the human hg38 reference genome using STAR (56) with default parameters.

Using BAM alignment, we tagged the location of each R1 by their barcode and position in the reference genome (54). Sequences ranging from 4 to 8 base pairs surrounding each insertion site were extracted to identify common octanucleotide sites listed in Supplementary Table 3.

#### Identification of DNA binder recognition sequences in the human genome

We used custom python scripts to search for octanucleotide target sequences in the human genome reference sequence (hg38) and generated a list of genomic coordinates parsed from an entire genome FASTA file. DNA binder (ZnF or TALE) recognition sequences occurring within the specified number of base pairs of the target site were compiled. The resulting sequences were aligned to the human genome to determine exact genomic coordinates.

### Genomic PCR

Lysates from the safe harbor genome targeting experiments using helper plasmids designed to target the chr 6 site (ALFA MLT m2, chr 6L NbALFA, and chr 6R NbALFA) were used as PCR template to amplify the junction between the transposon and genome. Similar to above, 1 µl of lysate, 10 µl of KOD ONE BLUE polymerase 2x mastermix (Toyobo), 7.8 µl water and 0.6 µl each primer was used. The forward primer extended from the transposon (Left end For or Right end For) and the reverse primer extended from the genome (chr 6 genome For). PCR cycle conditions were 95°C for 30 sec followed by 32 cycles of 98°C for 10 sec, 60°C for 5 sec, and 68°C for 10 sec. Products were run on a 2% agarose gel and cleaned using the Zymo Gel DNA Recovery Kit (Zymo). For sequencing, cleaned PCR products were ligated into pJET1.2/blunt sequencing vector using the CloneJET PCR Cloning Kit (Thermo Fisher Scientific). Primer sequences are listed in Supplementary Table 2.

## RESULTS

### Mutations in the native MLT DNA-binding domain reduce random genomic integration

To develop a conditional transposase for INTACT, we hypothesized that off-target insertions could be minimized by disrupting the interaction between the transposase’s native DNA-binding domain and genomic DNA. By subsequently introducing a DNA-binding protein, such as a programmable ZnF or TALE, the transposase could be directed to the target sequence, thereby restoring integration activity (Fig. 1a).

**Fig. 1.**
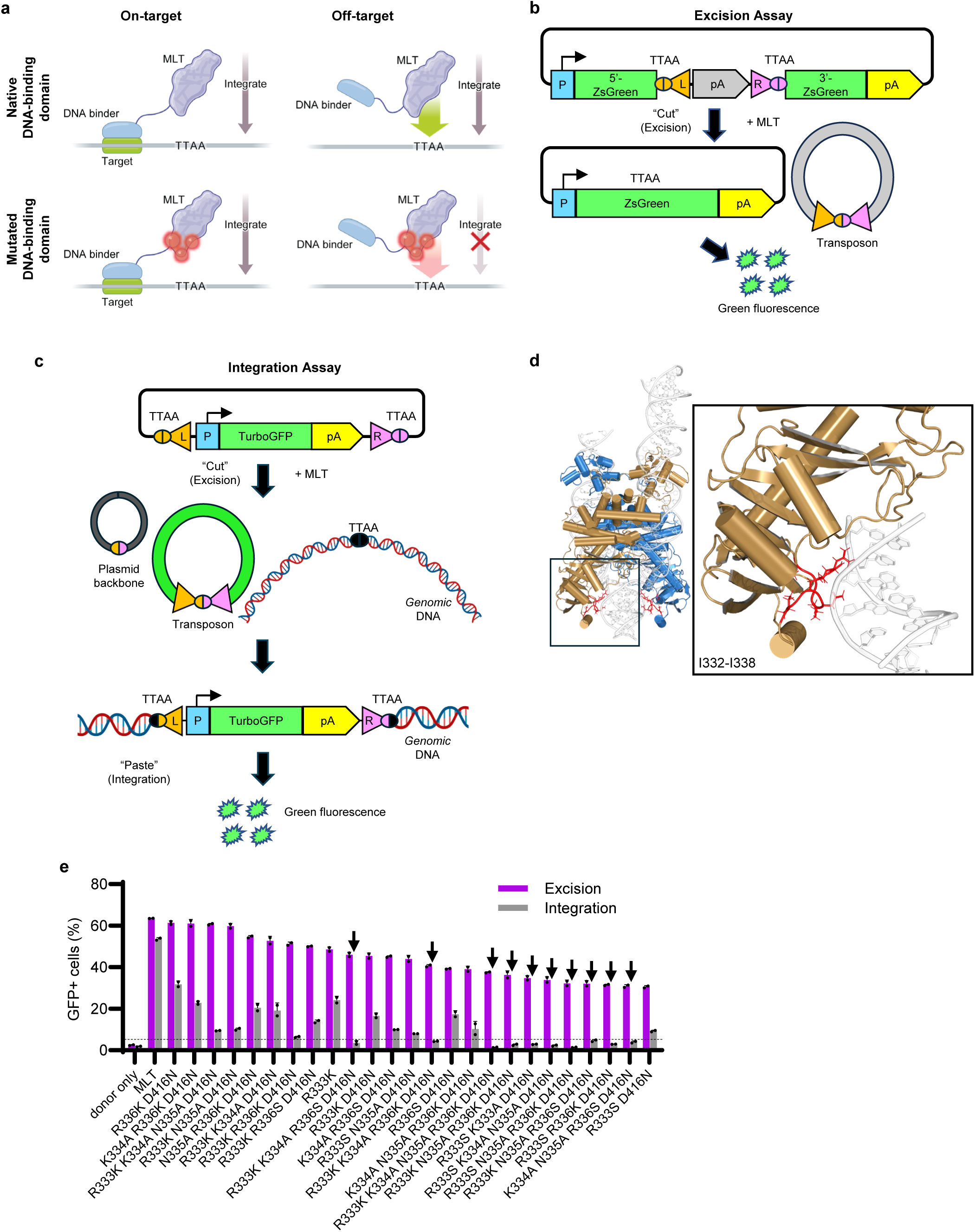
Identification of Exc^+^Int⁻ DNA-binding domain mutants to minimize off-target integration. **a**, Proposed model for improved specificity by disrupting the MLT DNA-binding domain. Both the native (top left panel) and mutant (bottom left panel) transposases integrate at on-target sites via exogenous DNA binder targeting. The native transposase retains full DNA-binding activity, allowing for off-target integration (top right panel). In contrast, the mutant transposase shows reduced off-target integration due to impaired native DNA-binding (bottom right panel). **b**, Successful excision of the transposon restores a complete ZsGreen reading frame. Excision activity is quantified by measuring ZsGreen fluorescence by flow cytometry. **c**, Stable TurboGFP integration is assessed by flow cytometry after 17 days, once transient expression from unintegrated plasmid is lost. **d**, AlphaFold2 structural model of MLT identifying amino acid residues (I332-I338) in the predicted DNA-binding domain (highlighted in red). **e**, MLT excision (purple bars) and integration (grey bars) across a panel of DNA-binding domain mutants. Ten excision-capable, integration-deficient mutants (Exc^+^Int⁻) (m1–m10, black arrows) were selected based on excision levels >30% and integration levels <5% (dotted line). *n* = 2. Data are shown as mean + s.d.

We used the structure of MLT predicted by AlphaFold2 aligned to the *piggyBac* strand transfer complex (*pdb 6×67*) using PyMOL (31,57) to identify regions of the protein that, when mutated, would impair its DNA-binding function and reduce integration while preserving its catalytic activity for excision (Exc⁺Int⁻). Although additional positively charged residues likely contribute to non-specific DNA interactions, structural data from *piggyBac* (31) indicate that the loop spanning residues 332-338 contains key determinants for DNA engagement. Guided by these structural insights and prior mutagenesis in *piggyBac* (39), we targeted this loop in MLT to generate Exc⁺Int⁻ variants. This strategy aims to weaken DNA binding sufficiently to reduce off-target integration while retaining residual contacts that allow integration to be restored when the transposase is localized by a DNA-binding protein.

In all experiments, we employed a previously characterized hyperactive transposase (Supplementary Fig. S1a and Supplementary Note). To screen for Exc⁺Int⁻ mutants, we designed excision and integration assays, both of which were quantified using flow cytometry. The excision assay uses a reporter gene disrupted by an MLT transposon (Fig. 1b). Upon expression of the transposase and seamless excision of the transposon, the reporter gene is reformed, resulting in expression. The integration assay uses a plasmid with a transposon containing a constitutively expressed reporter gene that is inserted into the genome. Cells with stable insertions are quantified following 17 days of culture (Fig. 1c).

To identify Exc⁺Int⁻ mutants, we performed alanine scanning of residues along the loop (I332-I338) predicted by AlphaFold2 to interact with DNA (Fig. 1d). Substitutions I332A, R333A, R336A, G337A, and I338A strongly impaired both integration and excision, while substitutions K334A and N335A slightly reduced these activities (Supplementary Fig. S1b).

Given that nonpolar, hydrophobic alanine substitutions appeared too disruptive to DNA binding, we next evaluated polar but uncharged serine, which can engage in hydrogen bonding with DNA, as a less disruptive alternative. We also tested positively charged lysine, which can form electrostatic interactions with DNA and potentially counterbalance the effects of alanine or serine when used in combination. Given the predicted importance of direct interactions of arginine residues R333 and R336 with DNA, we focused our panel of mutants to positions spanning R333-R336, with residues R333 and R336 either left unchanged or converted to lysine or serine. K334 and N335, shown to tolerate alanine substitution, were left unchanged or replaced with alanine. Because mutations in the putative DNA-interacting loop might reduce excision activity, we included the MLT D416N mutation due to its structural homology to the *piggyBac* mutant, D450N, known to improve excision in the context of integration-dampening mutations (39). We performed a pilot screen in duplicate of 64 combinations of mutations. In total, 10 mutations exhibited >30% excision efficiency and <5% integration efficiency (Fig. 1e and Supplementary Fig. S1c,d). Notably, all 10 contained the D416N mutation. One mutant, containing R333K, K334A, N335A, R336K and D416N (herein referred to as m1), exhibited a 1.4% integration efficiency, close to the background signal without transposase helper plasmid. Therefore, we chose m1 for further characterization.

### Linking artificial DNA-binding proteins to MLT targets integration to plasmids in human cells

In an effort to target our Exc⁺Int⁻ mutant to a specific sequence during INTACT, we linked the transposase to the well-characterized ZnF protein E2C, that binds the 9 bp recognition sequence called ‘e2c’ (58). We reasoned that the small size (∼90 amino acids) would offer optimal targeting efficiency while minimizing the risk of steric hindrance or interfering with transpososome formation.

We developed a plasmid-to-plasmid (P2P) targeting assay to screen transposase frameworks (Fig. 2a). The P2P assay uses a donor plasmid in which the transposon contains a promoter driving the first 5’-half of an EmGFP gene followed by a splice donor sequence. A reporter plasmid contains the target sequence followed by a splice acceptor and the 3’ fragment of EmGFP. Successful excision of the transposon followed by insertion at the target sequence results in restored EmGFP expression that is measured by flow cytometry (Fig. 2a). PCR primers are used to amplify the transposon-reporter junction to identify the location of insertions. In the initial design, we included a 640 bp target region containing 26 TTAA sites and the e2c recognition sequence to study targeting performance in a localized region of competing integration sites. A control reporter plasmid, containing the identical target region, but lacking the e2c recognition sequence, was used for measuring untargeted integration.

**Fig. 2.**
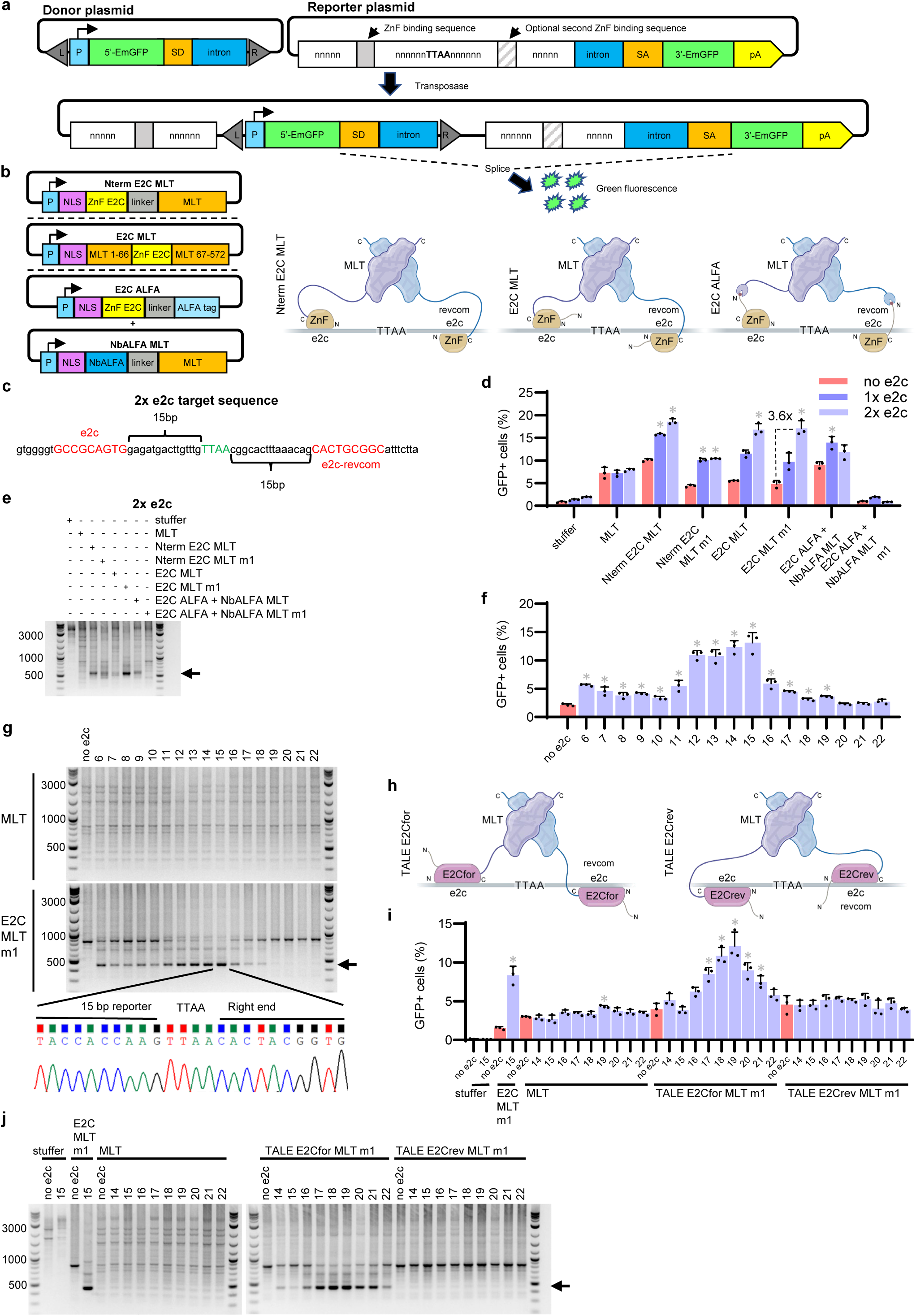
Linking MLT to DNA-binding proteins mediates targeting to plasmids. **a**, ZnF targeting of MLT to either one or two binding sites on the P2P reporter plasmid results in insertion of the transposon at the target TTAA. Splicing of the two halves of EmGFP leads to expression quantified by flow cytometry. **b**, Helper plasmids for distinct strategies linking the ZnF to MLT. The ZnF was covalently fused to the N-terminus (Nterm E2C MLT), embedded in the MLT protein (E2C MLT), or linked through a nanobody-tag association (E2C ALFA + NbALFA MLT). **c**, Target sequence of the 2x e2c reporter. **d**, MLT linked to the E2C ZnF mediates targeting of a transposon to reporter plasmids containing single (dark purple) or flanking (light purple) e2c recognition sites. Red bars represent reporters with no e2c recognition site. Unmutated MLT was compared to the Exc⁺Int⁻ mutant m1. Stuffer control was plasmid backbone without transposase. *n* = 3. **e**, PCR analysis of the P2P assay from **d**. Products indicated by the black arrow represent insertions that occurred near the target e2c sequence. **f**, Spacing constraints for E2C MLT m1. P2P reporter plasmids contained equidistant e2c recognition sequences, spaced 6 to 22 bp from the target TTAA on the left and right. *n* = 3. **g**, PCR analysis of the P2P assay from **i**. Products indicated by the black arrow represent targeted insertions. Representative Sanger sequencing chromatogram of the 15 bp reporter PCR product, displaying the reporter plasmid – transposon junction sequence. **h**, TALEs inserted after the G66 residue of MLT were designed to bind the e2c recognition sequence or its reverse complement. The recognition sequences were positioned opposite each other on the P2P reporter target region. **i**, Spacing constraints for TALE-based targeting of MLT. P2P reporter plasmids contained e2c recognition sequences, spaced 14 to 22 bp from the target TTAA on the left and right. Unmodified MLT was compared to MLT fused to TALEs (**h**). Stuffer control was plasmid backbone without transposase. *n* = 3. **j**, PCR analysis of the P2P assay from **i**. Products indicated by the black arrow represent targeted insertion ns. Data are shown as mean + s.d. (*) = P value < 0.01, derived from a Student’s two-tailed t-test.

Initial attempts at INTACT explored three transposase frameworks (Fig. 2b) including E2C fused to the transposase N-terminus (Nterm E2C MLT), E2C inserted after the glycine at position 66 within the N-terminus (E2C MLT), and E2C tethered to MLT through a nanobody bound to an ALFA-tag peptide (E2C ALFA + NbALFA MLT) (49).

For the second framework (E2C MLT), we hypothesized that the unstructured N-terminal domain may serve as a *de facto* ‘flexible linker’ for covalently linked DNA-binding proteins. By incorporating the ZnF within the N-terminal domain, we shortened this linker and positioned the catalytic domain closer to the target DNA. The third framework (E2C ALFA + NbALFA MLT) used a non-covalent strategy that connected the DNA-binding protein to the transposase via an interaction allowing for transient dissociation, potentially allowing them to separate and undergo necessary conformational changes during catalytic insertion.

Helper, donor, and reporter plasmids were co-transfected into HEK293T cells and lysed three days later. PCR analysis of the P2P integration products demonstrated that all three frameworks generated prominent products representing insertions near the e2c recognition sequence (Supplementary Fig. S2a). PCR products were sequenced revealing that insertions occurred at a TTAA 15 bp downstream of the e2c recognition sequence. Because MLT forms a dimer, we hypothesized that efficiency might improve with two e2c recognition sequences flanking a central TTAA (Fig. 2c) (29). Indeed, reporter plasmids with dual e2c recognition sites showed increased targeting efficiency by flow cytometry and brighter PCR products of the expected size (Fig. 2d,e and Supplementary Fig. S2b). The m1 mutation abolished activity in the non-covalent system, potentially due to suboptimal configuration when linked to the N-terminus. The m1 MLT Exc⁺Int⁻ mutant with an internal E2C DNA-binding protein at the G66 position (E2C MLT m1) produced 1.8-fold more GFP+ cells on the 2x e2c reporter plasmid relative to the 1x e2c reporter, and 3.6-fold more GFP+ cells compared to the reporter lacking an e2c recognition site (no e2c). (Fig. 2d). E2C MLT m1 displayed an especially prominent PCR product of the expected size and DNA sequence (Fig. 2e and Supplementary Fig. S2b).

To determine optimal spacing between the DNA recognition site and the target TTAA, we assayed reporter plasmids with e2c recognition sequences spaced 6 to 22 bp on either side of the target TTAA and removed all other TTAAs within a 100 bp window of the target site to reduce confounding effects. Consistent with previous results, the optimal spacing for E2C MLT m1 was between 12 and 15 bp, with 14 and 15 bp spacing having the highest signal (Fig. 2f,g).

To attempt INTACT at endogenous human sites, we identified loci with identical 9 bp ZnF recognition sequences spaced either 14 or 15 bp from a central TTAA on either side. We synthesized 29 ZnFs by modular assembly (59–62) (Supplementary Table 4) and inserted them after the G66 residue of the Exc⁺Int⁻ mutant MLT m1. Helper and donor plasmids were co-transfected into HEK293T cells, but genomic targeting was not detectable by ddPCR after 3 days (data not shown). This may be due to inherent difficulties in engineering high-affinity ZnF proteins to user-defined sequences (63,64). Therefore, we tested TALEs as alternate DNA-binding proteins.

We generated two TALEs, one to the forward and another to the reverse complement (revcom) of the e2c recognition sequence. Each TALE was inserted after the G66 residue in the Exc⁺Int⁻ mutant (TALE E2Cfor MLT m1 and TALE E2Crev MLT m1) (Fig. 2h). The P2P assay revealed optimal targeting at 18 or 19 bp for the forward TALE but no targeting using the reverse TALE (Fig. 2i,j). As with ZnFs, we observed that targeting was improved by flanking e2c recognition sites (Supplementary Fig. S2c,d). To evaluate an alternative target sequence to e2c, we tested TALEs designed to bind a sequence in the human *CCR5* gene using our P2P assay (33). By swapping in the CCR5 TALEs, we observed that TALEs matched to their corresponding recognition sequences supported targeting, while mismatched did not (Supplementary Fig. S2e,f). Based on these results, we continued our targeting experiments using TALEs in place of ZnFs.

### The target TTAA flanking sequence influences the insertion choice by Exc⁺Int⁻ mutants

In our P2P assays, we observed reproducible PCR products at off-target locations, which we attributed to the presence of preferential TTAA sites within the reporter plasmid. For example, we observed that mutant m1 consistently inserted the transposon at a specific TTAA (shown at 854 bp in gels from Fig. 2e,g,j and Supplementary Fig. S2a,b,d,f). Sequencing revealed that this specific TTAA was flanked by TA dinucleotides (taTTAAta) and integrations at this site appeared to compete with the target TTAA. TA pyrimidine-purine base steps are known to confer greater intrinsic DNA bending, leading us to hypothesize that MLT Exc⁺Int⁻ mutants, due to their impaired native DNA-binding, may prefer more deformable target TTAA sequences such as the taTTAAta octanucleotide (65). Interestingly, the related transposase *piggyBac* requires a tight bend in the target TTAA to facilitate integration (31).

We note that the P2P flow cytometry analysis detects both on-target insertions between the DNA-binding protein recognition sequences in addition to insertions occurring at nearby off-target sites such as the taTTAAta octanucleotide. To improve the sensitivity of our P2P assay by reducing competing signal from the preferred off-target taTTAAta site, we disrupted the TTAA by mutating the third thymine to a guanosine (taTGAAta) (Supplementary Fig. S3a). After validating the improved reporter lacking the off-target octanucleotide, we screened nine additional Exc⁺Int⁻ mutants (E2C MLT m2-m10) to identify variants with enhanced activity. Mutant m2 (R333K, N335A, R336K, D416N) showed a twofold increase in targeting efficiency compared to m1 (Fig. 3a and Supplementary Fig. S3b).

**Fig. 3.**
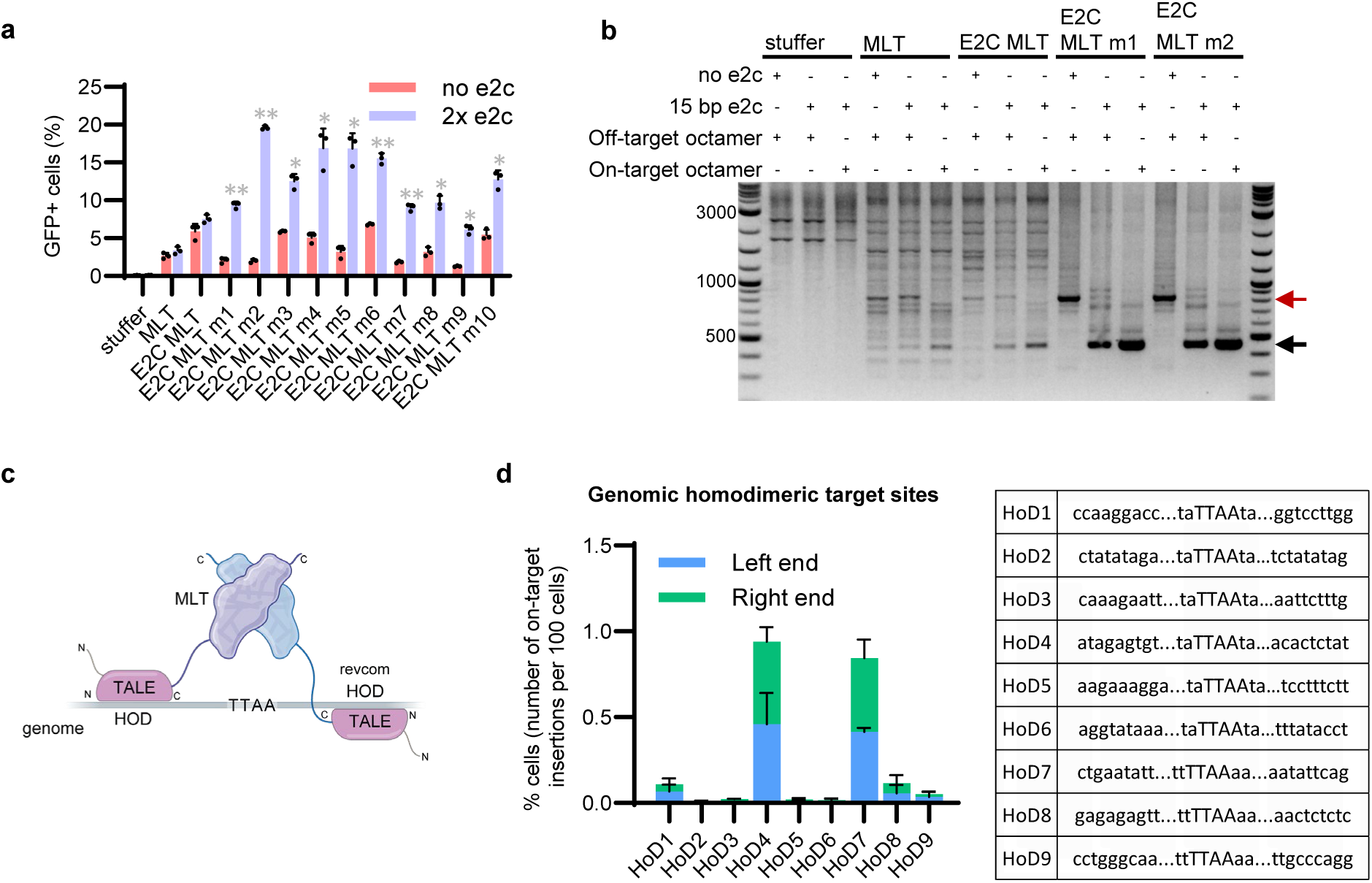
MLT targeting to endogenous sequences at preferred octanucleotide sites. **a**, Comparison of E2C ZnF targeted MLT variants with Exc^+^Int⁻ mutations in the I332-I338 loop, predicted to interact with DNA. P2P assay quantification of integration into reporter plasmids with 2x e2c recognition sites (light purple) or no e2c (red). MLT was unmodified without a ZnF. Stuffer control was plasmid backbone without transposase. *n* = 3. **b**, The taTTAAta octanucleotide is a preferential integration site for Exc⁺Int⁻ mutants linked to the E2C ZnF. PCR analysis of the P2P assay using reporter plasmids with the octanucleotide positioned off-target (upper red arrow) or on-target between the 2x e2c recognition sites (lower black arrow). Exc⁺Int⁻ mutants m1 and m2 were compared to fusions without mutations. MLT was unmodified without a DNA binding protein. Stuffer control was plasmid backbone without transposase. **c**, TALEs designed to bind homodimer (HoD) recognition sequences flanking target TTAA sites in the human genome were inserted after the G66 residue of the MLT mutant m2. **d**, TALEs targeting nine genomic homodimer sites were each designed to bind an identical 9 bp target sequence flanking either taTTAAta (HoD1–6) or ttTTAAaa (HoD7–9) octanucleotides located 18 or 19 bp on either side of the TTAA. ddPCR for measuring the per-cell on-target insertion efficiency was performed using a forward primer and probe in the genome and a reverse primer at either the left or right transposon end. The number of on-target insertions per 100 cells was determined by multiplying the target-to-reference ratio by the number of reference copies per cell. *n* = 3. Data are shown as mean + s.d. (*) = P value < 0.01, (**) = P value < 0.00005, derived from a Student’s two-tailed t-test.

To further validate the taTTAAta octanucleotide as a preferential integrating site, we modified the reporter plasmid to contain the octanucleotide flanked by e2c recognition sequences 15 bp apart. We observed a prominent targeted PCR product band using E2C MLT m2 on the taTTAAta reporter plasmid (Fig. 3b). Similar results were obtained using a TALE in place of the ZnF on reporter plasmids containing TALE binding sites 19 bp apart (Supplementary Fig. S3c). Subsequent experiments exploited the m2 mutant’s preferential targeting of taTTAAta octanucleotide sites.

### Covalent fusion of a TALE to MLT targets octanucleotide sites in the human genome

Given the observed preference for insertion at the taTTAAta octanucleotide site on our reporter plasmid, we investigated whether other octanucleotide sites were similarly favored. To address this, we analyzed a recently generated database of genomic insertion sites for unmodified MLT, obtained using a modified self-reporting transposon (SRT) protocol (see Methods) (54). The most frequently targeted octanucleotide was ttTTAAaa, while the previously identified taTTAAta site ranked 21st out of 136 sites (Supplementary Table 3).

To investigate INTACT at genomic octanucleotide sites, we identified loci where identical 9 bp sequences flanked a target octanucleotide site spaced 18 bp or 19 bp apart. Homodimeric (HoD) flanking sites were selected to enable targeting with a single TALE. We chose target sites located more than 50 kb from coding genes, yielding six taTTAAta sites (HoD1–6) and three ttTTAAaa sites (HoD7–9) (Supplementary Table 5). TALEs designed to bind the flanking recognition sequences of these sites were inserted after the G66 residue in the m2 mutant (Fig. 3c).

Genomic targeting efficiency was quantified by ddPCR, with the HoD4 site showing the highest on-target insertion rate at 0.9% of cells (Fig. 3d). We measured an approximately equal number of insertions from the left and right ends, indicating a lack of preference for insertion in either orientation. Control experiments confirmed that targeting was TALE-mediated rather than being driven by insertion preferences at octanucleotide sites (Supplementary Fig. S3d). Due to the aneuploid nature of HEK293T cells, which can contain variable numbers of chromosomes or chromosomal fragments, we used ddPCR to quantify available target sites. We measured an average of ∼8 HoD7 sites per cell, approximately threefold more than HoD4 (Supplementary Table 5). Despite similar integration rates for HoD4 and HoD7 per cell, the higher copy number of HoD7 corresponded to a lower per-target efficiency (0.4% for HoD4 compared to 0.1% for HoD7). This demonstrates that even when individual site-targeting efficiency is low, the overall per-cell integration rate can be enhanced by increasing the number of available target sequences. Together, these results suggest that TALEs direct transposon insertions at detectable levels in the human genome.

### Non-covalent linkage of TALEs to MLT improves targeting efficiency to the human genome

To improve the integration efficiency of INTACT, a non-covalent strategy was employed to allow greater flexibility during the integration reaction. The fusion of the DNA-binding domain directly to the transposase was hypothesized to constrain conformational changes required for integration (Fig. 4a), and transient separation of these components might relieve this constraint (Fig. 4b,c).

**Fig. 4.**
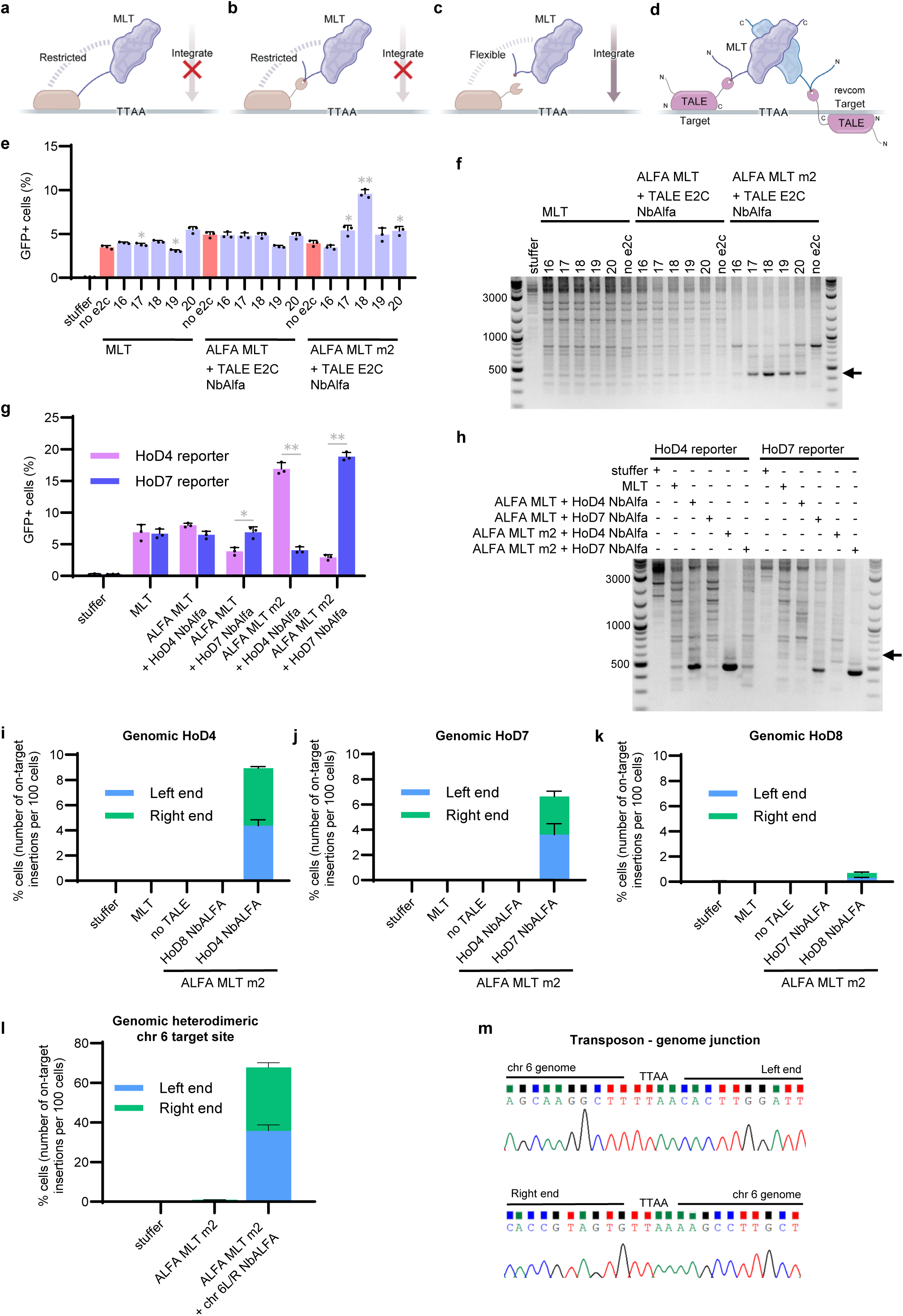
A non-covalent linkage of TALEs to MLT results in efficient targeting to the genome. **a**-**c**, Model for enhanced genome integration through transient disruption of the DNA binder-MLT linkage. Covalent linkage (**a**) or non-covalent linkage between the bound nanobody and ALFA tag (**b**) restricts MLT integration activity. Transient dissociation of the nanobody and ALFA tag (**c**) allows conformational flexibility, facilitating integration. **d**, ALFA MLT contained an ALFA tag inserted after the G66 residue of MLT. TALE NbALFA consisted of a genome-targeting TALE with a nanobody that recognizes the ALFA tag. Genomic targeting was enabled by the non-covalent interaction between MLT and the TALE via the nanobody-tag linkage. **e**, Spacing constraints for the non-covalent targeting approach using ALFA MLT and TALE NbALFA. P2P reporter plasmids contained e2c recognition sequences, spaced 16 to 22 bp from the target TTAA on the left and right. Exc⁺Int⁻ mutant m2 was compared to a fusion without mutations. MLT was unmodified without an ALFA tag. Stuffer control was plasmid backbone without transposase. *n* = 3. **f**, PCR analysis of the P2P assay from **e**. Products indicated by the black arrow represent targeted insertions. **g**, Non-covalent linkage of MLT to TALEs mediated targeting to genomic sequences on P2P reporter plasmids. Light purple bars indicate reporter plasmids with the HoD4 target site; dark purple bars indicate reporter plasmids with the HoD7 site. Matched and mismatched TALE-target site pairs were compared. *n* = 3. **h**, PCR analysis of the P2P assay from **g**. Products indicated by the black arrow represent targeted insertions. **i**-**k**, A non-covalent linkage strategy enables TALE-directed MLT insertion at genomic homodimeric sites. Controls included transfections without transposase helper plasmid (stuffer), unmodified MLT, ALFA MLT m2 without a TALE NbALFA fusion, and fusions targeting alternate genomic sites than the indicated target site (mismatched). *n* = 3. **l**, Heterodimeric TALEs direct integration at a genomic safe harbor site. ALFA MLT m2 was directed by different TALEs left (chr 6L NbALFA) and right (chr 6R NbALFA) of the target TTAA site on chr 6 and on-target insertion efficiency was measured by ddPCR. Controls included transfections without transposase helper plasmid (stuffer), and ALFA MLT m2 without a TALE NbALFA fusion. *n* = 3. **m**, DNA sequencing chromatograms depicting the left and right junctions of the transposon inserted at chr 6 from **l**. Data are shown as mean + s.d. (*) = P value < 0.01, (**) = P value < 0.00005, derived from a Student’s two-tailed t-test.

Our initial non-covalent strategy using the Exc⁺Int⁻ mutant m1 failed to promote targeted integration when the NbALFA nanobody was fused to the N-terminus of MLT (Fig. 2b,d). To improve performance, we adopted a revised configuration modeled after the successful TALE HoD MLT constructs: the 13-amino-acid ALFA tag was inserted internally after the G66 residue in the Exc⁺Int⁻ mutant m2 (ALFA MLT), and the nanobody was instead fused to the TALE (TALE NbALFA) to enable binding to the ALFA tag present on the transposase (Fig. 4d). The optimal spacing between the DNA-binding site and the target TTAA sequence was assessed using the P2P assay. The non-covalent framework maintained a spacing of 18 bp, comparable to the covalent strategy using the same TALE (Fig. 4e,f). We then generated P2P reporter plasmids containing the previously identified genomic homodimeric loci HoD4 and HoD7 target sites and observed increased targeting efficiency when TALEs were matched to their corresponding target sites compared to mismatched sites (Fig. 4g,h).

Genomic targeting to homodimeric loci using the non-covalent framework achieved an 8.9% per cell targeting efficiency at the HoD4 site, representing a >9-fold improvement over the covalent system (Fig. 4i). Targeting to the HoD7 site (6.6%) and HoD8 site (0.7%) also improved (Fig. 4j,k).

To further optimize INTACT, we swapped the NbALFA and ALFA tag between the TALE and transposase and inserted these domains after the S55 residue to increase linker length. We compared Exc⁺Int⁻ mutants m1 and m2 with unmodified MLT. However, none of these modifications outperformed the previous top-performing combination with ALFA tag inserted at G66, HoD4 NbALFA + ALFA MLT m2 (Supplementary Fig. S4a). Given the observed increase in efficiency of genomic targeting, the non-covalent approach was used in subsequent experiments.

### Heterodimeric TALEs mediate targeting to a genomic safe harbor

For purposes of simplicity, our initial INTACT optimizations used a single DNA-binding protein with identical homodimeric recognition sequences flanking the target TTAA. To expand the number of genomic loci available for integration, we explored the use of two TALEs designed to bind different heterodimeric sequences flanking the TTAA (Supplementary Fig. S4b). Taking advantage of this flexibility, we chose a target site on chromosome 6 (chr 6) with several desirable characteristics that was: (i) a preferred octanucleotide site (ttTTAAaa), (ii) a previously recovered MLT insertion site (from the SRT database described above), (iii) in open chromatin, (iv) in a genomic safe harbor site as defined by Papapetrou et al. (66), and (v) >120 kb from the nearest gene. We generated two TALE NbALFA fusions designed to recognition sequences spaced 18 bp on either side of the chr 6 TTAA site (chr 6L NbALFA and chr 6R NbALFA) (Supplementary Table 6). Co-transfecting these TALE fusions with ALFA MLT m2 resulted in a 67.8% per cell targeting efficiency by ddPCR (Fig. 4l) Genomic PCR confirmed the expected transposon-genome junction sequence (Fig. 4m). Furthermore, changing the TALE recognition sequences to bind shorter 8 bp or longer 10 bp and 11 bp sequences did not improve targeting compared to the 9 bp recognition sequences (Supplementary Fig. S4c). These data demonstrate that heterodimeric TALEs enable efficient genome targeting to a single safe harbor site.

### Optimizations of INTACT enhance genome targeting

The chr 6 site is located at a single genomic locus. To enhance INTACT integration efficiency, we hypothesized that increasing the number of potential target sites would be beneficial. A recent study demonstrated the use of precise RNA-mediated insertion of transgenes (PRINT) for targeted gene delivery to rDNA loci (67). The rDNA arrays exist in hundreds of copies per genome, providing sufficient redundancy such that insertional disruption of individual units does not compromise cellular function, thereby supporting their classification as a safe harbor (67,68).

Furthermore, rDNA arrays remain stable during mitosis in normal human somatic tissues, and prior studies have demonstrated successful expression of therapeutic proteins from rDNA-integrated transgenes in both mice and human cells, including blood clotting factors such as Factor VIII and IX (Hemophilia A and B), fumarylacetoacetate hydrolase (deficient in tyrosinemia type I), and mini-dystrophin (69–73). However, a limitation of the PRINT system is that the integration machinery often terminates prematurely, preventing full transgene insertion. In contrast, the “cut and paste” mechanism of transposases (74) ensures complete DNA cargo insertion during INTACT.

To determine whether INTACT could target rDNA, we generated heterodimeric TALEs for three sites within rDNA intergenic spacer regions, designated rDNA1, rDNA2, and rDNA3. For each site, TALE NbALFA fusions were designed to recognize sequences spaced 18 bp apart on either side of the TTAA site. rDNA1 and rDNA3 were both previously identified octanucleotides (rDNA1: taTTAAta, rDNA3: ttTTAAaa) and rDNA2 was a new octanucleotide that contained flanking ‘tt’ and ‘ta’ sequences and was the third most common octanucleotide used by MLT (Supplementary Table 3) (rDNA2: ttTTAAta). Both rDNA1 and rDNA2 were previously recovered MLT insertion sites from the SRT database. Two sets of TALEs were designed per site: one recognizing 9 bp sequences and the other recognizing 11 bp sequences (Supplementary Table 6). Both the 9 bp and 11 bp configurations for rDNA1, as well as the 11 bp set for rDNA3, outperformed our previous chr 6 safe harbor targeting strategy (Fig. 5a). We observed variability in integration, likely due to sensitivity to variations in day-to-day transfection efficiency, including a reduction in efficiency for chr 6 targeting compared to earlier experiments. Despite this variability, within the same transfection, rDNA targeting achieved a two-fold improvement over the chr 6 site, with a >56% per cell targeting efficiency for the three best conditions.

**Fig. 5.**
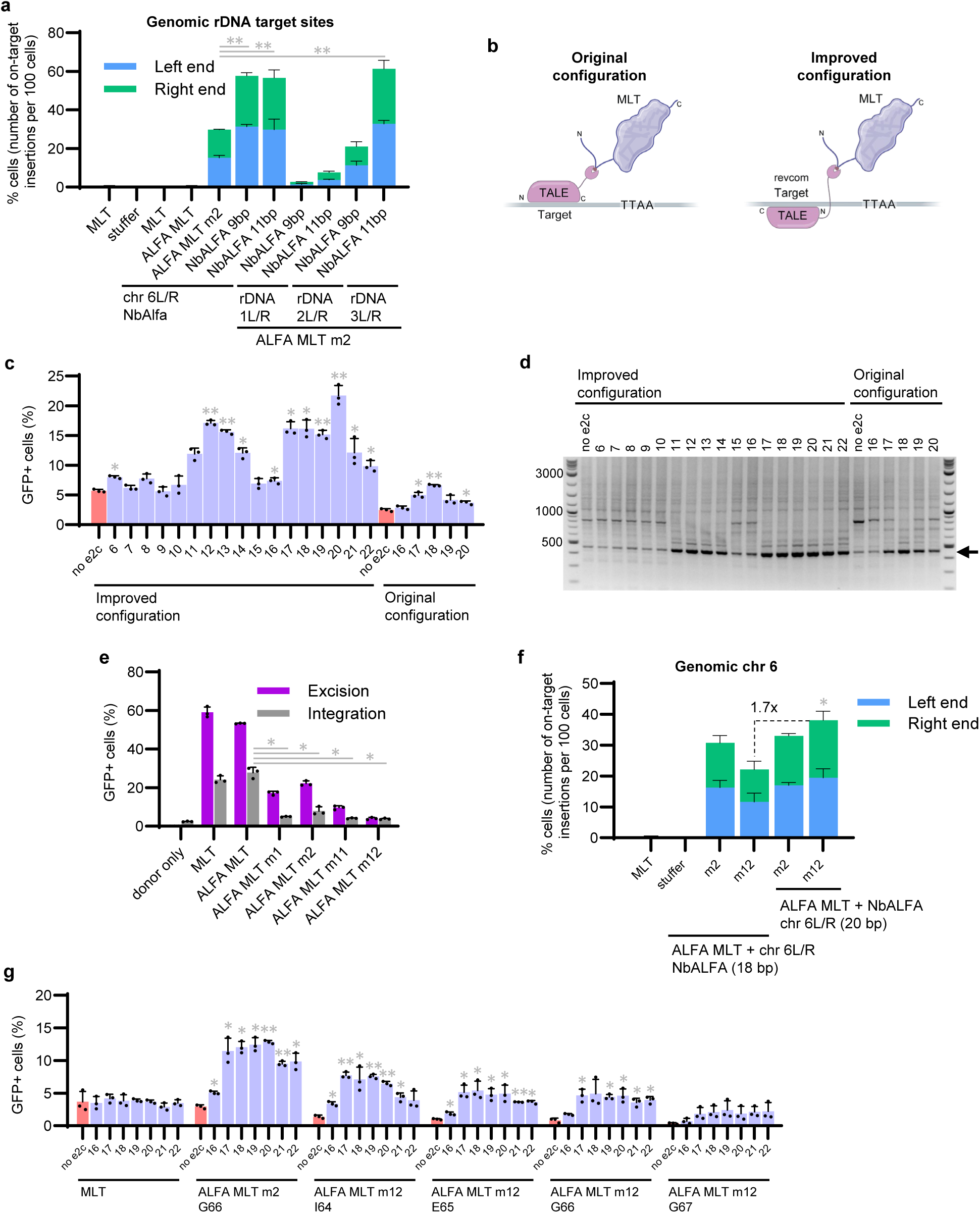
A panel of optimizations contributes to improved genome targeting. **a**, ALFA MLT m2 was directed by flanking left and right (L/R) TALEs designed to bind three rDNA target sites (rDNA1, rDNA2, and rDNA3). Both 9 bp and 11 bp TALE recognition sequences were tested. For comparison, chr 6 TALE NbALFA fusions were paired with ALFA MLT without an Exc⁺Int⁻ mutation or ALFA MLT m2. Controls included TALE NbALFA fusions without transposase (stuffer), and MLT without an ALFA tag that was with or without a TALE NbALFA fusion. *n* = 3. **b**, To create an improved configuration, the TALE was redesigned to bind the reverse complement of the original recognition sequence, and the NbALFA nanobody was repositioned from the C-terminus to the N-terminus. **c**, Spacing constraints for the improved configuration NbALFA TALE E2Crev using a TALE that binds the reverse complement of the e2c recognition sequence. P2P reporter plasmids contained e2c recognition sequences, spaced 6 to 22 bp from the target TTAA on the left and right. *n* = 3. **d**, PCR analysis of the P2P assay from **c**. Products indicated by the black arrow represent targeted insertions. **e**, MLT excision (purple) and integration (grey) comparing ALFA MLT Exc^+^Int⁻ mutants m1, m2, m11, and m12. Controls included ALFA MLT without an Exc^+^Int⁻ mutation and MLT without an ALFA tag. Donor only control did not include a helper plasmid and represents background non-enzymatic inserted plasmid. *n* = 3. **f**, ALFA MLT mutants m2 or m12 were directed by heterodimeric TALEs flanking the target TTAA site on chr 6 and on-target insertion efficiency was measured by ddPCR. Two fusion configurations were tested: NbALFA fused to the N-terminus (NbALFA TALE) or to the C-terminus (TALE NbALFA) of the TALE. *n* = 3. **g**, Spacing constraints for ALFA MLT m12 were assessed by varying the position of the ALFA tag (I64–G67) within the unstructured N-terminal domain. ALFA MLT m12 was paired with the NbALFA TALE E2Crev fusion designed to bind to the reverse complement of the e2c recognition sequence. P2P reporter plasmids contained e2c recognition sequences, spaced 16 to 22 bp from the target TTAA on the left and right. ALFA MLT m2 with ALFA tag at G66 and MLT without an ALFA tag were included for comparison. *n* = 3. Data are shown as mean + s.d. (*) = P value < 0.01, (**) = P value < 0.00005, derived from a Student’s two-tailed t-test.

We next investigated the orientation of the TALE NbALFA fusion, hypothesizing that relocating the NbALFA from the C-terminus to the N-terminus of the TALE and redesigning the TALE recognition site to the reverse complement (revcom) sequence would position NbALFA similarly in space to our current configuration with the NbALFA at the C-terminus (Fig. 5b). To test whether this new configuration improved targeting, we employed the E2C P2P assay, which reports efficiency over a range of base-pair spacings. Using a TALE designed to bind the reverse complement of e2c (TALE E2Crev) with an NbALFA nanobody fused to the N-terminus (NbALFA TALE E2Crev), we observed a three-fold increase in targeting efficiency over the previous configuration, with an optimal spacing shift from 18 bp to 20 bp (Fig. 5c,d). We also observed local efficiency maxima at 12 bp. We speculate that the two observed peaks at 12 and 20 bp might be due to the twist of the DNA and the positioning of the TTAA in relation to the catalytic site.

When identifying Exc⁺Int⁻ mutants to disrupt the native MLT DNA-binding domain, we initially selected mutants with >30% excision efficiency and <5% integration efficiency (Fig. 1e and Supplementary Fig. S1c). Given the high efficiency of our latest version of INTACT, we reevaluated mutants with lower excision rates (>15%). Two additional mutants, m11 (R333S, K334A, R336K, D416N) and m12 (R333S, K334A, N335A, R336K, D416N), which exhibited lower excision rates but near-background integration levels, were tested for potential reductions in off-target integration compared to m2. To measure off-target activity, we re-performed excision and integration assays, now with the ALFA tag insert, for ALFA MLT m1, m2, m11, m12 and without Exc⁺Int⁻ mutation. Donor without transposase was included to estimate random background plasmid insertion. Our previous mutant m2 displayed a clear increase in integration over background, while m12 exhibited near-background integration levels (Fig. 5e). Consequently, we selected m12 for future configurations.

Encouraged by the improvements seen in the P2P assay for NbALFA nanobody fusions at the N-terminus of TALE E2Crev, we generated NbALFA TALE fusions targeting the chr 6 safe harbor site, incorporating a 20 bp spacing from the TTAA in the reverse complement orientation relative to our prior configuration (NbALFA chr 6L and NbALFA chr 6R). These fusions were paired with ALFA MLT m2 from prior experiments or ALFA MLT m12 which reduced off-target integration to near-background levels. The highest on-target insertion levels were observed for the NbALFA TALE with the m12 mutation, a 1.7-fold improvement over the prior configuration with m12 (Fig. 5f).

Until this point, we had only tested ALFA tag fusions at residues S55 and G66. The S55 fusion, featuring a longer linker, was predicted to increase flexibility between the DNA-binding protein and transposase but resulted in lower on-target genomic insertion levels than G66 (Supplementary Fig. S4a). To assess how ALFA tag positioning could affect targeting levels, we inserted the ALFA tag at adjacent residues (I64, E65, and G67). Since repositioning the ALFA tag could affect optimal spacing, we used the P2P assay with NbALFA TALE E2Crev with a known preference for 20 bp spacing when the ALFA tag was inserted at G66 and reporter plasmids containing spacing of 16 bp to 22 bp to capture any changes from the new positioning. The m2 mutant with G66 fusion yielded higher on-target levels but also increased off-target activity, as indicated by signal from the negative reporter plasmid lacking e2c binding sites (Fig. 5g and Supplementary Fig. S5a). Notably, m12 with ALFA tag fusion at I64 resulted in higher on-target insertion than G66, albeit with a slight increase in off-target levels. Fusion at G67 significantly reduced both on– and off-target activity, suggesting that this linker length was too short. Based on these findings, we retained G66 as the optimal residue for ALFA tag fusion due to its minimal off-target activity and sufficient flexibility without sacrificing overall efficiency.

### High efficiency genome targeting to rDNA safe harbor sites

Having determined the optimal configuration for fusion of the NbALFA domain to the N-terminus of the TALE, 20 bp spacing between the TALE and TTAA, the m12 mutation for reduced off-target integration, and ALFA tag fusion at G66, we applied these optimizations to rDNA1 (octanucleotide taTTAAta) (Fig. 6a). Both upstream and downstream TALEs were designed, ranging from 12 bp to 17 bp in length (Supplementary Table 6). Systematic testing of all possible combinations revealed that a 12 bp downstream TALE paired with a 15 bp upstream TALE achieved the highest per cell targeting efficiency of 49.9% (Fig. 6b). An added benefit of this configuration is that longer 12 bp and 15 bp TALE recognition sequences occur less frequently in the genome than the 9 bp sequences flanking the chr 6 site, potentially enhancing specificity.

**Fig. 6.**
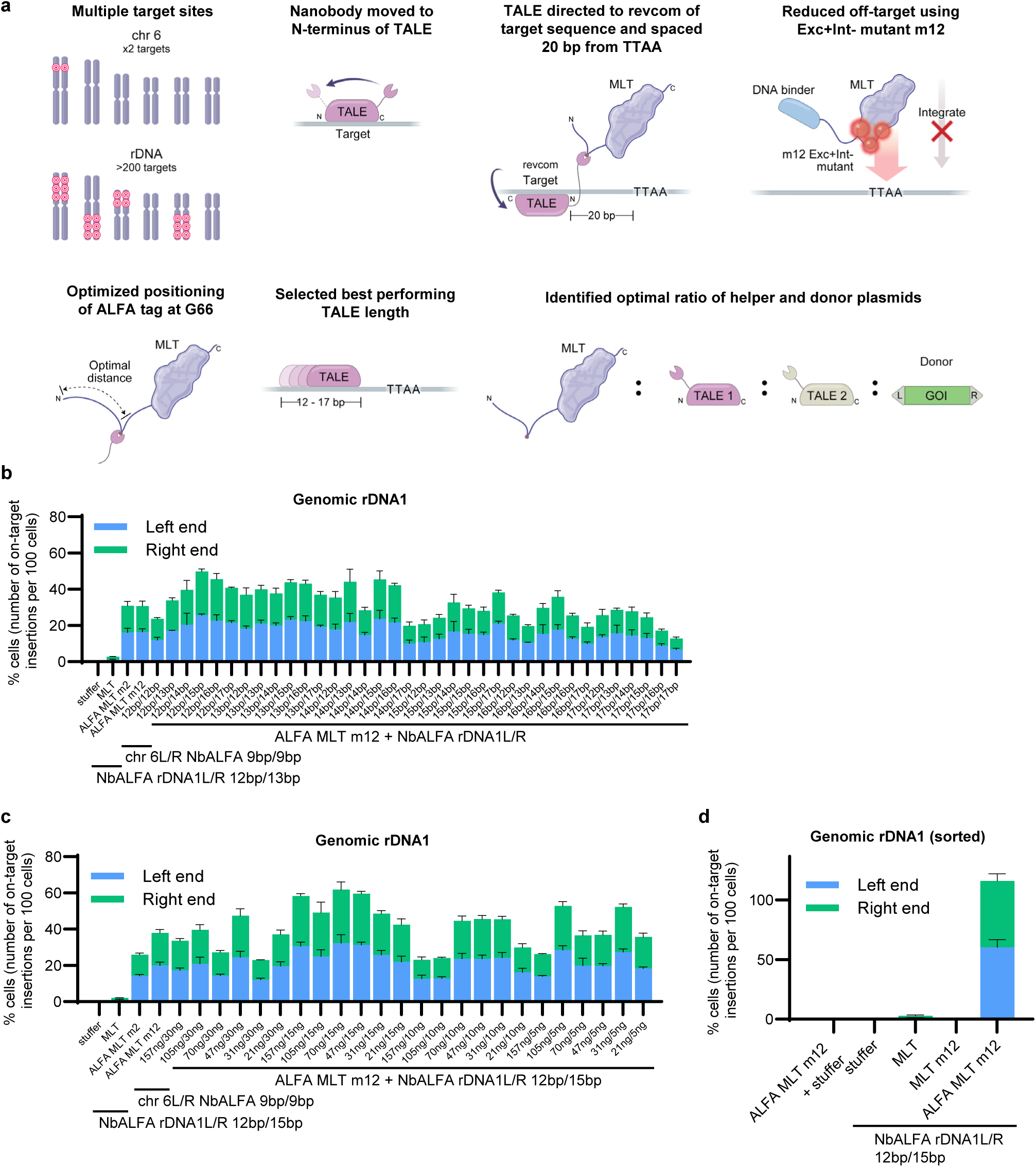
High efficiency targeting to multi-copy rDNA sites. **a**, A series of optimizations enhanced INTACT genome targeting efficiency, including: increasing the number of available target sites by targeting rDNA; repositioning the NbALFA nanobody to the TALE N-terminus; reversing TALE orientation to bind the reverse complement of the recognition sequence, spaced 20 bp from the target TTAA; using the Exc⁺Int⁻ m12 mutant to reduce off-target insertions; optimizing ALFA tag positioning within the unstructured N-terminal domain of MLT; varying TALE recognition sequence length; and tuning the helper-to-donor plasmid ratio. **b**, ALFA MLT m12 was directed to the rDNA1 site by flanking left and right (L/R) TALEs designed to bind recognition sequences ranging from 12 bp to 17 bp in length. On-target insertion efficiency for all combinations of TALE lengths was measured by ddPCR. For comparison, ALFA MLT mutants m2 and m12 were paired with chr 6 TALE NbALFA fusions. Controls included rDNA1 NbALFA TALE fusions without transposase (stuffer) or with unmodified MLT. *n* = 3. **c**, Integration efficiency was optimized by varying helper-to-donor plasmid ratio. The ng of helper over donor plasmid is indicated. *n* = 3. **d,** On-target integration efficiency at rDNA1 incorporating a panel of optimizations. Cells were sorted for iRFP670 fluorescence to ensure successful transfection and DNA uptake. Controls included ALFA MLT m12 without TALEs, NbALFA TALEs without transposase helper plasmid, unmodified MLT with NbALFA TALEs, and MLT m12 without an ALFA tag and with NbALFA TALEs. *n* = 4. Data are shown as mean + s.d.

For consistency across experiments, all transfections were performed with 70 ng of transposase-encoding MLT helper plasmid, 15 ng of each TALE-expressing plasmid, and 500 ng of donor plasmid. To evaluate the impact of plasmid stoichiometry on integration efficiency, we varied the amounts of ALFA MLT m12 helper plasmid (21-157 ng) and rDNA1 TALE plasmids (5-30 ng). Integration efficiencies across the 24 conditions ranged from 22.9% to 61.8%, with the original helper-to-donor ratio yielding the highest efficiency (Fig. 6c).

To control for variations in transfection efficiency during INTACT, we performed fluorescence-based sorting to enrich for cells strongly expressing a co-delivered plasmid and exclude poorly transfected cells. We also extended the culture period from three to four days to allow additional time for MLT expression. We included an additional control of ALFA MLT m12 without NbALFA TALE fusions. Also, to demonstrate that the nanobody-tag interaction was necessary, we generated a control helper plasmid by removing the ALFA tag from ALFA MLT m12 called MLT m12. By targeting multi-copy rDNA sites, it is possible to have more than one targeted insertion per cell. Sorting for successfully transfected cells improved efficiency, resulting in most cells receiving at least one targeted insertion, with an average of 1.2 on-target insertions per cell (Fig. 6d). Collectively, our panel of optimizations were additive (Fig. 6a), contributing to the development of an improved targetable transposase system.

## DISCUSSION

In this study, we developed INTACT, a mammalian transposase system that enables highly efficient, site-directed integration of gene-sized DNA sequences into genomic safe harbors. By leveraging rationally designed mutations within the native DNA-binding domain, we reduced untargeted integration while restoring insertion activity at desired sites through the addition of programmable TALE DNA-binding proteins.

A key innovation of our approach is the non-covalent linkage of the TALE via a nanobody-tag interaction. We suggest that this linkage permits conformational flexibility for formation and action of the transposome during the integration process. A similar principle has previously been used to non-covalently link the *Sleeping Beauty* transposase to a DNA binding protein. By fusing a domain called N-57, known to interact with *Sleeping Beauty*, to the TetR DNA binding protein, Ivics et al. successfully targeted pre-inserted TetR binding sequences in the genome (44).

We showed that linkage of TALE DNA-binding proteins within the flexible N-terminal domain of the transposase enhanced targeting efficiency. While previous transposase engineering efforts have primarily relied on direct N-or C-terminal fusions, our results indicate that linking the DNA-binding protein within the disordered N-terminal region preserves catalytic function while allowing for improved targeting.

To develop an efficient and specific targeting system, we conducted a large-scale screen to identify excision-positive, integration-negative (Exc⁺Int⁻) MLT mutants. Using structural predictions, we introduced targeted mutations to disrupt native DNA-binding while preserving excision activity. Key residues believed to be involved with DNA binding were originally identified by alanine-scanning mutagenesis in the *piggyBac* transposase (39). In the Alphafold2 model of MLT, this corresponding region on a loop spanning residues 332-338 was returned with >90% confidence and closely aligned with the cryo-EM structure of *piggyBac* (31). In this alignment, the amino acids targeted for mutagenesis are positioned in close proximity to the bound DNA (Fig. 1d). While we recognize that structural predictions cannot substitute for experimentally determined structures, the strong model confidence, combined with prior mutagenesis data, supports the interpretation that this loop contributes directly to DNA engagement. Consistent with this, substitutions within this region generated Exc⁺Int⁻ variants, highlighting its functional importance.

We used a dual-reporter system to measure excision and integration independently, allowing us to systematically select MLT mutants that retained excision activity but exhibited minimal integration. This screening strategy was essential for establishing a transposase architecture in which genomic integration is contingent upon co-delivery of a sequence-specific DNA-binding protein. By minimizing integration at off-target loci and restoring activity selectively at desired sites, our system provides a framework for programmable, high-specificity genome insertion.

Our findings reveal a strong preference for specific octanucleotide sequences (taTTAAta and ttTTAAaa), suggesting that local DNA bending may facilitate integration (65). This insight enabled us to identify and prioritize genomic sites that enhance targeting efficiency. We determined that the distance between the DNA-binding protein and the TTAA insertion site was a crucial factor in optimizing targeted integration. To precisely define the optimal spacing of INTACT, we developed plasmid-based assays that systematically varied the position of recognition sequences relative to TTAA sites. These assays revealed that integration efficiency peaked at a specific spacing, supporting the hypothesis that correct positioning of the transposase relative to its target sequence is essential for efficient DNA insertion.

Through a progression of optimizations, we demonstrated that targeted integration can be systematically enhanced through iterative refinement of transposase engineering, DNA-binding protein configuration, and genomic site selection. Unlike CRISPR-Cas systems, the recognition sequence for TALEs can easily be varied by adding or deleting binding repeats, which allowed us to fine-tune binding affinity by adjusting TALE length on either side of the target TTAA. Using TALEs targeting multi-copy rDNA loci, we attained targeting efficiencies as high as 1.2 on-target insertions per cell, reinforcing the potential of rDNA as a promising target for therapeutic applications requiring high efficiency. This represents a significant advancement in targeted transposition, surpassing the efficiencies reported for several previously engineered transposase systems (32–44).

While this manuscript was under review, an enhanced CAST system was reported that employed phage-assisted continuous evolution (PACE) to engineer a targetable transposase with substantially improved integration efficiency and no requirement for the cytotoxic bacterial accessory protein ClpX (75). This advancement underscores the distinct yet complementary trajectories of CAST and INTACT systems. CAST derives from bacterial systems, where the principal challenge has been to boost efficiency in mammalian contexts. In contrast, INTACT is based on the mammalian-derived MLT transposase, which inherently exhibits high activity in mammalian cells. Our focus was to enhance specificity and confine this high-efficiency integration to defined genomic loci.

INTACT offers several advantages over competing genome-editing technologies. Unlike methods that rely on twin prime editing, our method does not induce a pair of DNA nicks, thereby avoiding associated indel formation (22,76). Additionally, our system requires only two DNA-binding proteins and a single transposase, a substantial simplification compared to the 7-8 proteins and guide RNA required by CAST systems (47,75).

We used an unbiased integration assay to quantify off-target insertion into the genome. The MLT m12 mutation, used for high efficiency rDNA targeting, resulted in off-target insertion rates near background levels. Although future studies could be designed to map all off-target loci, our primary objective was to minimize the total frequency of off-target integrations. This is particularly relevant for chimeric antigen receptor (CAR) T-cell therapies, where conventional transposase-based methods lack targeting specificity (77). These therapies could benefit from targeted transgene insertion into rDNA loci shown to support high expression (67–73) while reducing the risk of insertional mutagenesis.

Because MLT functions as a dimer, the current heterodimeric TALE-transposase fusion strategy necessarily generates a mixture of homodimeric (TALE1-TALE1, TALE2-TALE2) and heterodimeric (TALE1-TALE2, TALE2-TALE1) species. Only the heterodimeric configuration optimally contributes to targeted insertion, whereas the homodimeric forms may interfere by binding transposon ends or occupying genomic TALE recognition sites. This distribution is reminiscent of early zinc finger nuclease designs, where half of the assembled species were similarly non-productive. In that case, rational engineering of the dimerization interface led to obligate heterodimers, achieved by introducing complementary charge residues that disfavored homodimerization while stabilizing heterodimer formation (78,79). A similar approach could be applied to INTACT to bias assembly toward productive heterodimers and thereby increase the fraction of complexes capable of targeted transposition. Implementing such a system would require additional orthogonal DNA binding protein linking strategies, for example, employing a second nanobody-epitope tag pair to ensure that each protomer associates with the intended TALE.

Future studies could explore additional strategies to enhance on-target integration efficiency, including testing a broader range of Exc⁺Int⁻ mutants or targeting alternative genomic loci. Precision could also be improved through protein-based delivery of preassembled transposase-transposon complexes, which may limit expression time and reduce potential off-target insertions (80). Previous studies have demonstrated that rDNA targeting can be enhanced by localizing the transposase to the nucleolus (81,82). This approach might complement rDNA-directed TALEs used here. Directed evolution offers another promising strategy to enhance transposase activity and specificity (53). To validate the therapeutic potential of INTACT, future studies are warranted for evaluating delivery of clinically relevant genes, such as for *ex vivo* CAR T-cell therapy or *in vivo* correction of monogenic disorders such as hemophilia (77,83).

In summary, this study establishes a novel site-directed transposase system that enables efficient, targeted integration of gene-sized DNA sequences without introducing DNA breaks. These findings provide a robust platform for transposase-based genome engineering, with broad implications for gene therapy, synthetic biology, and cell-based therapeutics.

## DATA AVAILABILITY

The data underlying this article are available in the article and in its online supplementary material.

## AUTHOR CONTRIBUTIONS

*Conceptualization*: J.B.O., J.F.M., J.J.H., and R.T. contributed to the initial project design and formulation of key hypotheses.

*Methodology*: J.B.O., J.F.M., B.E.H. and J.J.H. developed the experimental strategies and protocols used throughout the study.

*Investigation*: J.B.O., J.E.S. L.S., I.S., C.T.T., C.E., B.E.J., D.F.W., and S.X. conducted experiments and collected data.

*Visualization*: J.B.O., and J.E.S. prepared figures and graphical representations of the data.

*Funding acquisition*: J.B.O., J.J.H., and R.T. secured the financial support necessary to carry out the research.

*Project administration*: J.B.O. and J.J.H. coordinated and managed the overall progress of the project.

*Supervision*: J.B.O. and J.J.H. oversaw the research and provided scientific guidance throughout the project.

*Writing – original draft*: J.B.O. and J.J.H. wrote the initial version of the manuscript.

*Writing – review & editing*: J.B.O. and J.E.S. revised and edited the manuscript for intellectual content and clarity.

## ACKNOWLEDGEMENTS

We thank all members of the Owens Laboratory for their valuable discussions and support. We are especially grateful to Nancy L. Craig for insightful and thought-provoking conversations on the mechanisms of targeted transposition. We thank the Molecular and Cellular Immunology Core at the University of Hawaii for technical support with flow cytometry. Bioinformatic analyses were conducted by Watershed Bioinformatics in Cambridge, MA, under a contract from SalioGen Therapeutics.

## Supplementary Materials

### Supplementary Note

#### Hyperactive MLT

An MLT mutant (S8P, C13R) with higher excision and integration activity than wildtype (Supplementary Fig. S1a) was developed by SalioGen Therapeutics (1). This enzyme was the starting point for additional mutations aimed at reducing off-target integration (Exc^+^Int^-^). A potential explanation for why this mutant is hyperactive is that the S8P, C13R mutations may lead to a loss of the first putative casein kinase II phosphorylation site. It was previously shown that phosphorylation of the N-terminal domain of the related transposase, *piggyBac*, decreased transposition and that alanine substitutions increased transposition (2). Therefore, a loss of phosphorylation might be expected to increase transposition activity of MLT.

---

**Supplementary Fig. S1.**
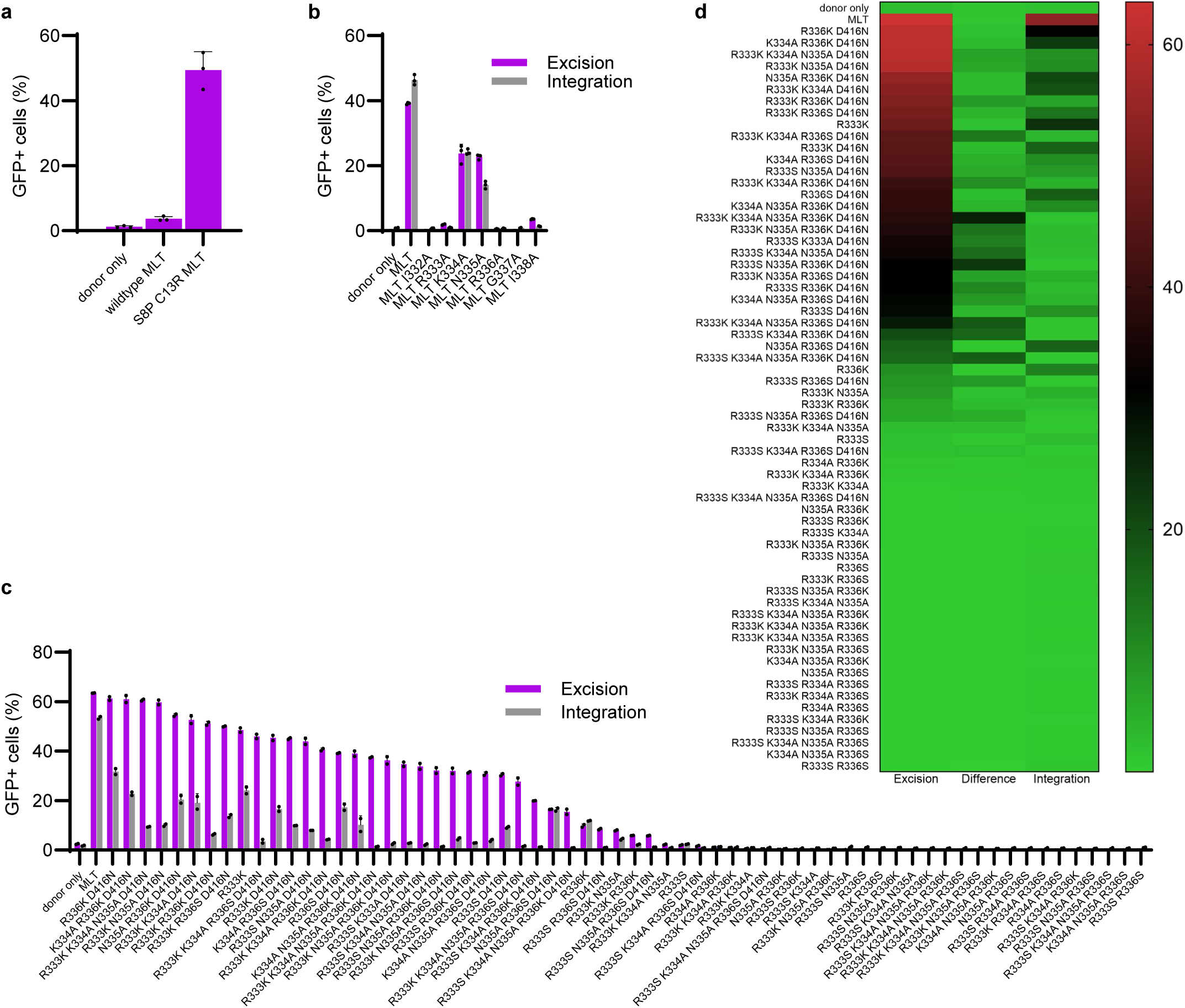
| Identification of DNA-binding domain mutants with reduced off-target integration. **a**, Comparison of integration efficiency of wildtype MLT and the hyperactive version of MLT with mutations S8P, C13R that was used in this study. *n* = 3. **b**, Alanine scanning of the predicted MLT DNA-binding domain (I332-I338). *n* = 3. **c**, Activities of 64 combinations of MLT DNA-binding domain mutations. Controls included MLT without mutations and ‘donor only’ without MLT helper plasmid. *n* = 2. **d**, Heatmap representation of the data from **c** of percentage of GFP+ cells depicting the difference in excision over integration in the center column. Left column excision, right column integration. Data are shown as mean + s.d.

**Supplementary Fig. S2.**
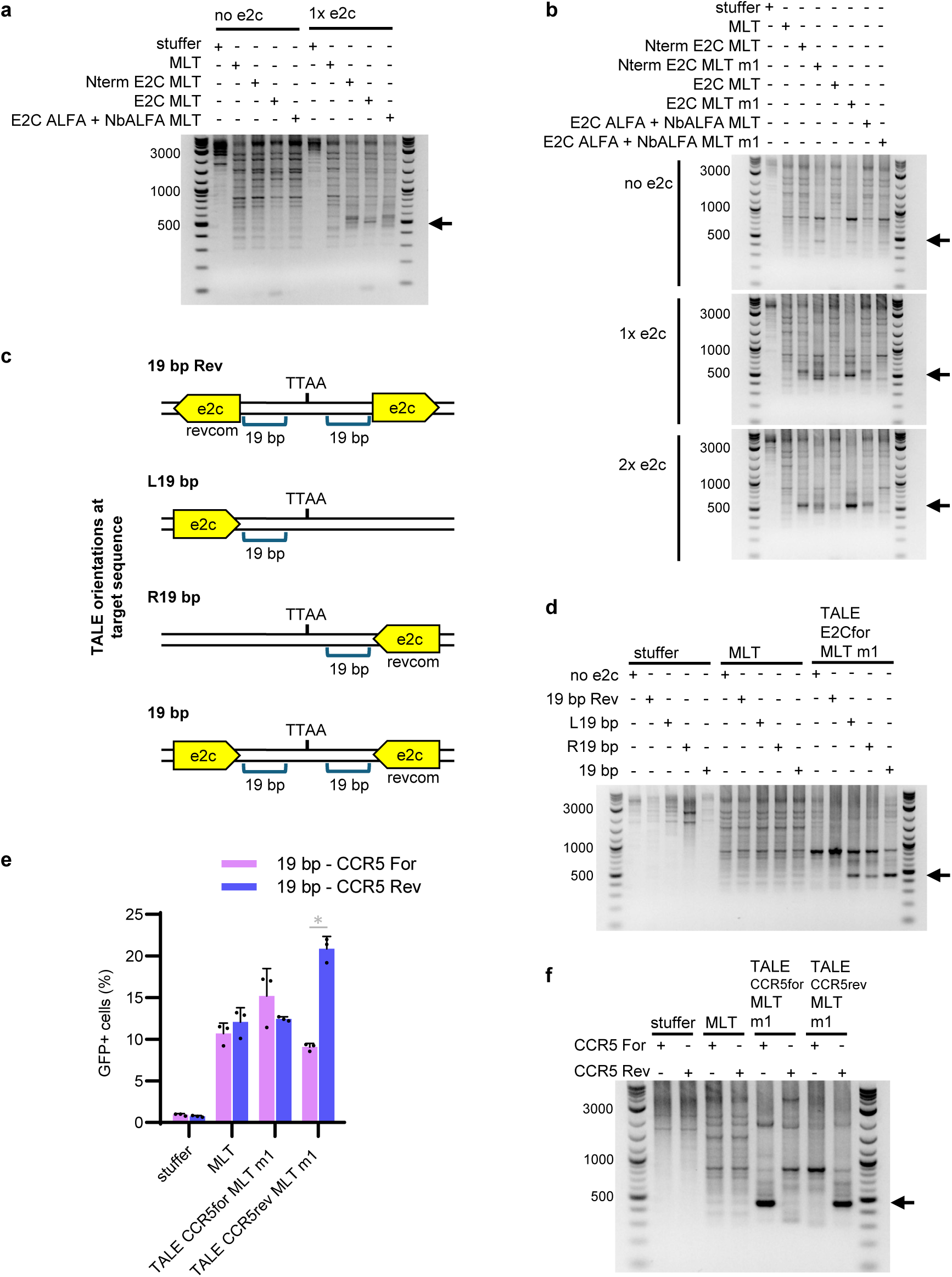
| Fusion of MLT to TALE or ZnF domains directs plasmid integration in human cells. **a**, PCR analysis of the P2P targeting assay for different strategies linking E2C to MLT. Products indicated by the black arrow represent insertions that occurred near the single target e2c sequence. **b**, PCR analysis of reporter plasmids containing zero, one, or two e2c recognition sites, using helper plasmids with or without the Exc⁺Int⁻ mutant m1. Products indicated by the black arrow represent targeted insertions. **c**, Reporter plasmids contain one or two e2c recognition sequences spaced 19 bp from the target TTAA. Yellow arrows indicate the direction of the e2c recognition sequences. **d**, PCR analysis of reporter plasmids from **c** containing one or two e2c recognition sites, using the MLT mutant m1 fused to an e2c-directed TALE. Controls included MLT without a TALE fusion or without MLT (stuffer). Products indicated by the black arrow represent targeted insertions. **e**, MLT fused to CCR5-directed TALEs was used to target reporter plasmids containing CCR5 recognition sites in either the matched or mismatched orientation. *n* = 3. **f**, PCR analysis of the P2P assay from **e**. Products indicated by the black arrow represent targeted insertions. Data are shown as mean + s.d. (*) = P value < 0.01 derived from a Student’s two-tailed t-test.

**Supplementary Fig. S3.**
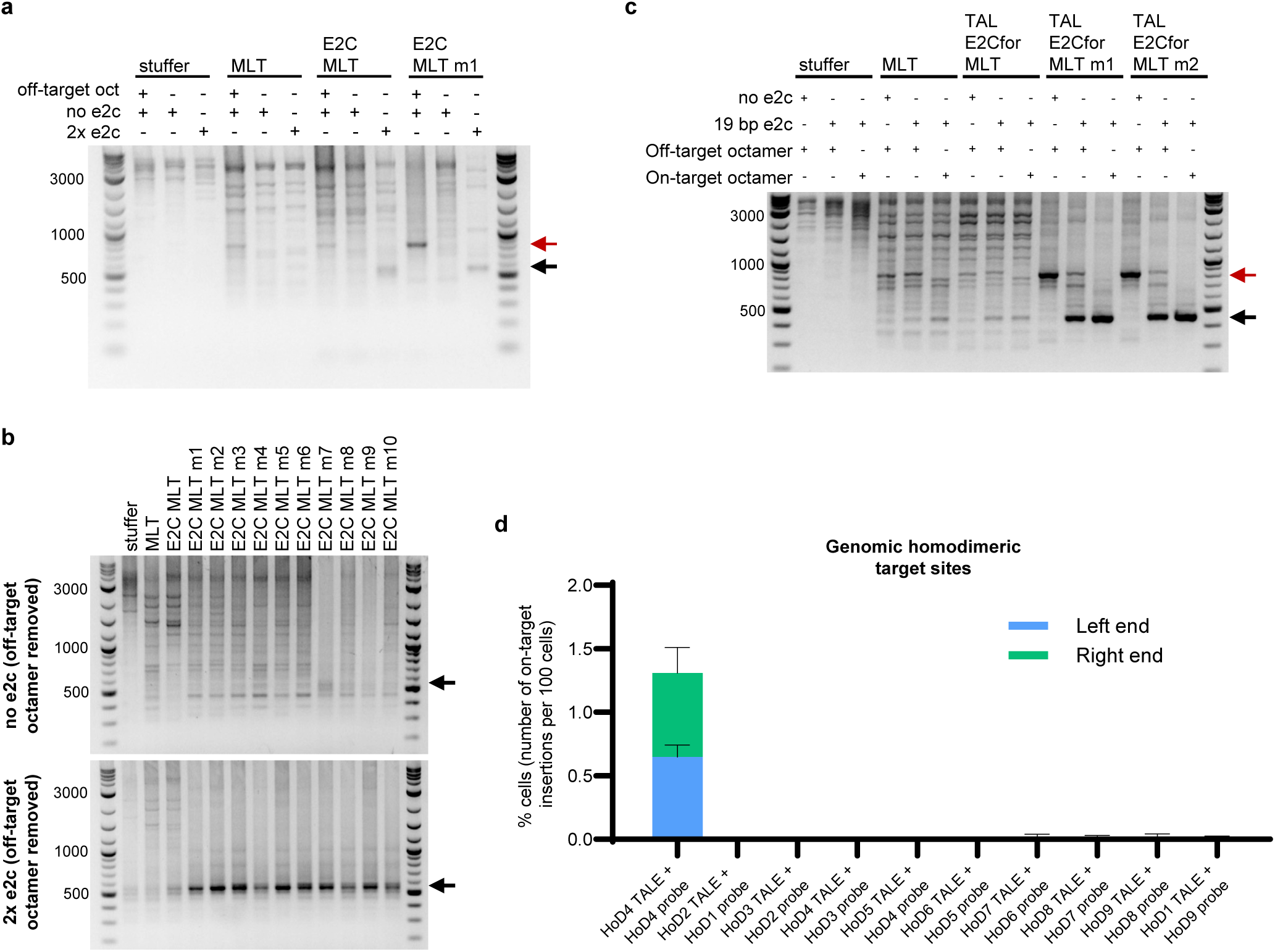
| A MLT DNA-binding domain mutant targets plasmids and endogenous genomic sequences. **a**, PCR analysis of reporter plasmids containing zero or two e2c target sites. To reduce competition between on-target and off-target TTAA sites within the plasmids, the off-target taTTAAta octanucleotide was mutated. The upper red arrow indicates the position of the off-target product, and the lower black arrow indicates targeted product. **b**, PCR analysis of the P2P assay using helper plasmids with Exc⁺Int⁻ mutations in the I332–I338 loop implicated in DNA binding. Reporter plasmids had a mutated off-target octanucleotide site (taTGAAta). Products indicated by the black arrow represent targeted insertions. **c**, The taTTAAta octanucleotide is a preferred integration site for Exc⁺Int⁻ mutants of MLT fused to the E2C TALE. PCR analysis of the P2P assay using reporter plasmids containing the octanucleotide either in an off-target location (upper red arrow) or positioned on-target between two E2C-binding sites (lower black arrow). Exc⁺Int⁻ mutants (m1 and m2) were compared to corresponding fusions lacking mutations. The unmodified MLT, lacking a DNA-binding domain, served as a control. Stuffer control was plasmid backbone without transposase. **d**, Matched and mismatched TALE-probe combinations were tested with helper plasmids targeting endogenous homodimeric genomic sites (HoD1–9). Mismatched combinations indicate background integration at TTAA sites. ddPCR was performed using a forward primer and probe in the genome and a reverse primer on the left end or right end of the transposon. *n* = 3. Data are shown as mean + s.d.

**Supplementary Fig. S4.**
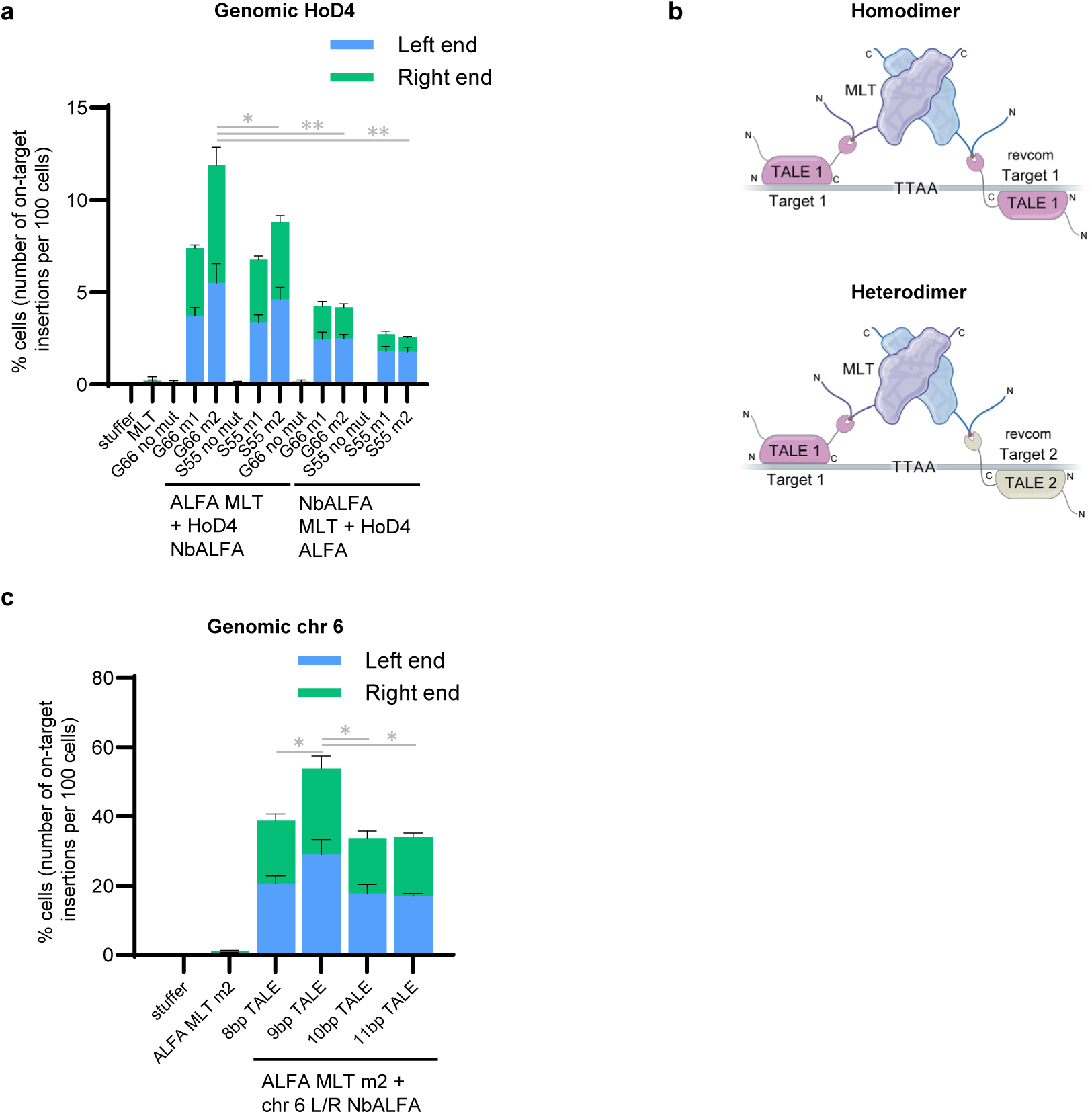
| Non-covalent tethering of TALEs to MLT enables efficient genomic targeting. **a**, Optimization of genome targeting to the HoD4 locus by adjusting ALFA tag and NbALFA nanobody positioning on MLT with or without Exc⁺Int⁻ mutations (m1, m2). The ALFA tag (ALFA MLT) or NbALFA nanobody (NbALFA MLT) were inserted after either G66 or S55 in the unstructured domain of MLT. The DNA binding component of the non-covalent system included the NbALFA nanobody (HoD4 NbALFA) or ALFA tag (HoD4 ALFA) fused to the C-terminus of the HoD4 TALE. *n* = 3. **b**, Targeting to a safe harbor site using heterodimeric TALEs. Unlike homodimeric TALEs, which require identical recognition sequences flanking the TTAA site, heterodimeric TALEs bind distinct sequences on either side, expanding the range of targetable loci. **c**, Comparison of TALEs designed to bind 8-11 bp of the chr 6 target sequence. ALFA MLT m2 was used either alone or with TALE NbALFA fusions recognizing the indicated base-pair lengths on the left and right sides of the target TTAA. *n* = 3. Data are shown as mean + s.d. (*) = P value < 0.01, (**) = P value < 0.00005, derived from a Student’s two-tailed t-test.

**Supplementary Fig. S5.**
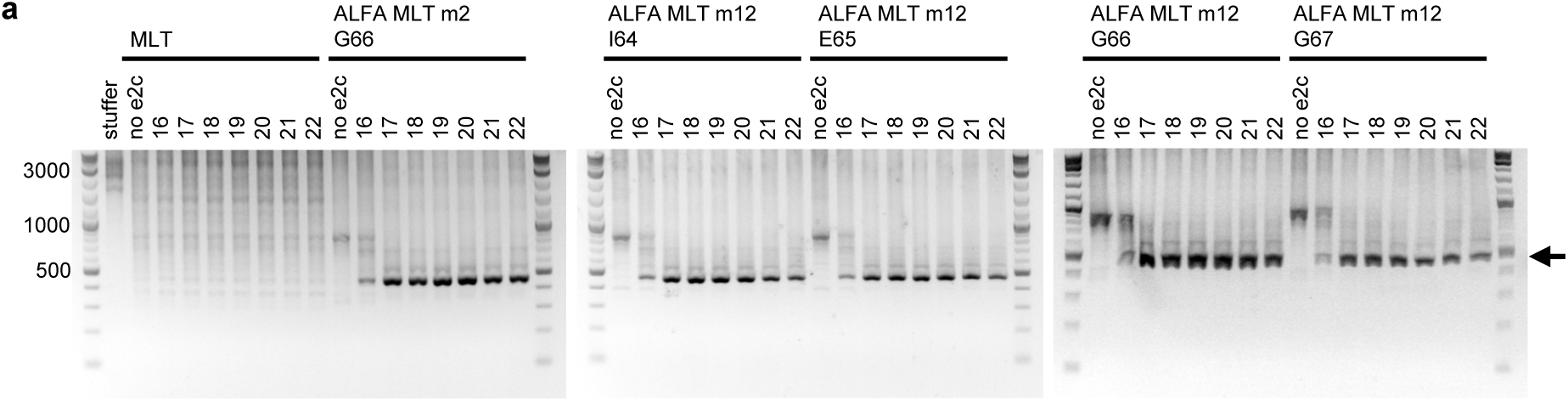
| Spacing constraints for ALFA MLT m12 were assessed by shifting the positioning of the ALFA tag within the N-terminal domain. **a**, PCR analysis of the P2P assay shown in Fig. 5h. P2P reporters contained e2c sites spaced 16-22 bp from the target TTAA on both the left and right and were co-transfected with helper plasmids containing the ALFA tag at the indicated positions (I64–G67). The black arrow indicates targeted insertions.

